# Transthyretin Contributes to Hippocampal Blood-Brain Barrier Recovery Following Intestinal Inflammation, with Reduced Expression in Alzheimer’s Disease

**DOI:** 10.64898/2026.09.01.748473

**Authors:** Zhi Xu, Jun Chen, Jiayi Yu, Jing Duan, Yazhou Xie, Yuqi Gong, Pan Wang, Xia Lei, Can Zhang, Xiaoduo Zhao, Zhongqin Fan, Bin Xu, Jing Zhang

## Abstract

**Background:** The blood-brain barrier (BBB) is increasingly recognized as an active immunoregulatory interface that responds dynamically to systemic inflammation. As intestinal inflammation can influence brain homeostasis through the gut-brain axis, the endogenous mechanisms that preserve BBB integrity during gut-derived inflammatory stress remain poorly understood.

**Methods and Results:** We combined dextran sulfate sodium-induced colitis, fecal microbiota transplantation, aged mice, APP/PS1 mice, human Alzheimer’s disease (AD) brain tissue, single-cell RNA sequencing, *in vivo* BBB permeability assays, and gain- and loss-of- function approaches to investigate adaptive neurovascular responses to gut inflammation and their mechanism in maintaining BBB integrity. Intestinal inflammation induced a region- specific adaptive response characterized by increased hippocampal vascular remodeling and partial restoration of BBB integrity following the initial inflammatory insult. Single-cell transcriptomic analysis identified a transthyretin (TTR)-enriched vascular-associated microglial state accompanying these neurovascular changes. Functional studies demonstrated that TTR contributes to maintaining BBB integrity by promoting endothelial homeostasis and limiting endothelial endocytosis, consistent with modulation of FcRn-associated transport pathways. This adaptive neurovascular response was progressively attenuated in aged mice and APP/PS1 mice and was accompanied by reduced vascular TTR expression in human AD brains.

**Conclusion:** These findings identify TTR as a contributor of adaptive neurovascular homeostasis during gut-derived neuroinflammation. Impairment of this homeostatic response with aging and AD may contribute to persistent BBB dysfunction and chronic neuroinflammation, highlighting neurovascular resilience as a potential therapeutic target.

## Introduction

While the blood-brain barrier (BBB) has traditionally been viewed as a static physical barrier separating the circulation from the brain, increasing evidence indicates that it is a dynamic interface that responds to systemic inflammatory stimuli.[1][2]. Dysregulation of BBB function is increasingly implicated in neuroinflammation and neurodegenerative disorders[3] [4], yet how BBB integrity is restored following peripheral inflammatory challenge remains poorly understood.

Among peripheral inflammatory stimuli, the gut-brain axis has emerged as a major regulator of central nervous system homeostasis[5]. Alterations in intestinal immunity and microbial composition influence brain function through circulating cytokines, microbial metabolites, extracellular vesicles, and immune cells, thereby modulating microglial activation, neuronal function, and vascular integrity[6–8]. Although chronic intestinal inflammation has been implicated in aging and Alzheimer’s disease (AD), the temporal response of the BBB to intestinal inflammation, including its capacity for subsequent recovery, remains incompletely understood.

The neurovascular unit consists of endothelial cells, pericytes, astrocytes, microglia, and neurons that collectively regulate BBB permeability and maintain the immune privilege of the CNS[9]. Recent studies have emphasized that BBB permeability is governed not only by tight junctions but also by tightly regulated transcellular transport pathways[10, 11], including receptor-mediated transcytosis. However, relatively little is known about the endogenous factors associated with BBB recovery following systemic inflammation, particularly in the hippocampus, a region highly vulnerable to aging and AD.[12].

Transthyretin (TTR) is best known as a carrier protein for thyroxine and retinol[13] and has long been implicated in AD through interactions with amyloid-β[14]. More recently, accumulating evidence suggests that TTR also influences endothelial function and vascular remodeling[15], raising the possibility that it may participate in regulating BBB homeostasis. Whether TTR contributes to the restoration of BBB integrity following peripheral inflammation, however, remains unknown.

Here, we demonstrate that intestinal inflammation induces a dynamic, region-specific adaptive response in the hippocampal BBB that is accompanied by increased TTR expression within the neurovascular unit. Functional studies identify TTR as a critical regulator of BBB integrity through modulation of endothelial function and FcRn-associated endocytosis. Importantly, this adaptive mechanism becomes progressively impaired in aged mice and is further diminished in AD. Together, these findings identify TTR as a previously unrecognized mediator linking gut-derived inflammation to neurovascular homeostasis and suggest that loss of this adaptive response may contribute to increased vulnerability of the aging brain.

## Methods and Materials

### Sex as a biological variable

This study included both male and female human participants. Sex was not considered as a biological variable in the analysis, and no sex-specific effects were examined. All animal experiments were conducted exclusively in male mice to improve experimental consistency. It remains to be determined whether the findings observed in male mice are fully applicable to female mice.

### Materials

The postmortem brain tissues were obtained from the China National Health and Disease Human Brain Tissue Resource Center (Hangzhou, China). A summary of the demographics and clinical data of the participants used for immunostaining is provided in **Table S1**. AD participants underwent extensive clinical evaluation and met specified inclusion and exclusion criteria. Details of the sample collection process have been previously described [16]. Wild-type (WT) mice (Stock No. SM-001, C57BL/6J) were obtained from Shanghai Model Organisms Center, Inc. APP/PS1 mice (Stock No. 005864, C57BL/6J background) were purchased from the Jackson Laboratory. The mice were housed under controlled conditions with a 12-hour light/dark cycle and unrestricted access to food and water.

### DSS-induced intestinal inflammation

Newly purchased 2-month-old WT (2m-WT) and 10-month-old WT mice (10m-WT) were randomized into cages with five mice each and housed for one week to normalize their gut microbiome. Intestinal inflammation was induced by administration of 3% DSS in drinking water for 1 day (T1) and 3 days (T3), followed by 2 days of regular water (T5), as described in a previous study [17].

### BBB permeability assay

BBB permeability was determined *in vivo* as previously reported [18, 19]. For the IgG extravasation assay, perfused mouse brain sections were stained with anti-mouse IgG1 antibody **(Table S2)**. Besides, 0.5% (w/v) Evans blue (EVB; #SL7203, Coolaber) in PBS was injected intravenously into the tail vein of mice on the indicated time point at a dosage of 10 μL/g. One hour after injection, mice were anesthetized and perfused with PBS. Brains and livers were excised and placed for optical imaging. The optical imaging system (OPTIMA, Biospace) was used to quantify the extent of EVB dye fluorescence accumulation with the excitation and emission filters set at 655 and 716 nm, respectively.

### Immunofluorescence staining

Mice were transcardially perfused with PBS under deep anesthesia. Brain and colon tissues were dissected, fixed in 4% paraformaldehyde, dehydrated in 30% sucrose, and sectioned by a cryostat (Leica). Sections were then incubated sequentially with a blocking solution (2% BSA, 0.4% Triton X-100, and 4% goat serum in PBS) for an hour at 26°C, primary antibodies overnight at 4°C, and corresponding secondary antibodies for an hour at 26°C **(Table S2)**. Z-stack confocal images were acquired using a FV3000 confocal microscope (Olmypus) and STELLARIS 8 confocal microscope (Leica).

### Single-cell RNA sequencing and data analysis

From T0 to T3, 2m-WT treated with DSS were sacrificed and perfused with PBS. Bilateral hippocampi were rapidly dissected and pooled together (n=3). Each group was replicated twice. We performed whole-cell droplet-based scRNA-seq from dissociated hippocampal tissue without nuclear isolation. It is well recognized that enzymatic dissociation of adult brain tissue leads to underrepresentation of mature neurons due to their large size and high sensitivity to mechanical and enzymatic stress. Sequencing was performed using the 10x Genomics Chromium platform (10x Genomics, Pleasanton, CA, USA) following the manufacturer’s protocol. Briefly, single-cell suspensions were loaded onto the Chromium controller to generate gel bead-in-emulsions (GEMs) for barcoding and reverse transcription. Libraries were prepared according to the standard 10x Genomics workflow and sequenced on an Illumina platform (NovaSeq 6000) to generate paired-end reads. Further downstream analyses were performed using the Seurat (v4) pipeline based on the function SCTransform. Clusters were determined using the function FindClusters with a resolution parameter of 0.2. Marker genes for each cluster were defined with the function FindAllMarkers with parameters min.pct = 0.25 and test.use = wilcox. Differential gene expression analysis for each cell type was performed across time points using the function FindMarkers with parameters min.pct = 0.20, avglogFC = 0.20, and test.use = wilcox. Intercellular ligand- receptor interactions were examined with the function CellChat analysis.

### Western blot (WB)

Tissues and cells were grinded (Jing xin), and the protein concentration was determined through the Protein Quantification Kit (#23225, Thermo Fisher Scientific). Samples were loaded onto gradient gels (GenScript) and transferred onto a nitrocellulose blotting membrane (#T500361, PALL). After blocking, the membranes were incubated with the primary antibody **(Table S2)** followed by incubation with an HRP-conjugated secondary antibody or IRDye® 800CW conjugated secondary antibody. Images were visualized using a ChemiDoc MP imaging system (Bio-Rad). The results were analyzed using ImageJ software with β-actin as internal controls.

### Hippocampus stereotaxic injection

WT were anesthetized and placed on the stereotaxic injection apparatus (RWD Life Science). After the skin was incised and the skull drilled, the mice were unilaterally injected with 0.1 μL pAAV5-CBh-mTtr-EGFP, pAAV5-U6-mTtr[shRNA]-mCherry or their corresponding controls (VectorBuilder) on the right side of the hippocampus (from bregma, anterior posterior: −1.7 mm; mediolateral: - 1.4 mm, dorsal ventral: −2.0 mm) using a microliter syringe (Hamilton) in 1 min. The needle was removed 5 minutes after the completion of the infusion. 2m-WT injected with pAAV5-CBh-mTtr-EGFP or control were maintained for 6 weeks and then administered intraperitoneally with LPS (#L4516, Sigma) at the dosage of 10 mg/kg 24 hours prior to optical imaging, to induce systemic inflammation, thereby modeling GBA dysfunction. 2m-WT injected with pAAV5-U6-mTtr[shRNA]-mCherry or control were housed for 6 weeks until optical imaging. 6-month-old WT mice (6m-WT) injected with pAAV5-CBh-mTtr-EGFP or control were housed for 4 months until behavioral test.

### Cell culture and treatment

hCMEC/D3, HMC3 and HBVP cells were obtained from ATCC and cultured in DMEM (Gibco) containing 10% FBS (Gibco) and 1% penicillin/streptomycin (Corning) and maintained at 37°C in 5% CO_2_ atmosphere. HMC3 and hCMEC/D3 cells were incubated with human TTR (hTTR) overexpressing lentivirus and negative lentivirus vectors (VectorBuilder) for 24 hours and then screened with puromycin at a dosage of 5 μg/mL. Medium from hTTR-overexpressing HMC3 and control were collected and centrifuged to obtain the supernatant for further studies.

### Quantitative real-time PCR (qPCR)

The qPCR analysis was performed according to the instructions (#Q711, Vazyme). The primers for quantification of human *Ttr* mRNA are CGTGCATGTGTTCAGAAAGGCTG (Forward) and CTCCTCAGTTGTGAGCCCATGC (Reverse).

### Cell migration assay

TTR overexpressing hCMEC/D3 cells and controls were seeded into each well of the Ibidi Culture-Insert 2 Well system (#80206, Ibidi), which were pre-positioned in a 48-well plate. The culture inserts were carefully removed when cells reach 100% confluence. Cells were rinsed with medium and covered with fresh growth medium. Pictures of migrating cells were taken and evaluated at 0 h and 24 h after removal of inserts.

### Tube formation assay

Matrigel was thawed on ice, diluted by pre-chilled culture medium and then evenly spread onto a pre-chilled 96-well plate at 100 µL per well, ensuring it covers the entire bottom surface. The plate was incubated at 37℃ for 1 h to allow the Matrigel to solidify. hCMEC/D3 cells were carefully seeded onto the solidified matrigel in each well at the density of 2 × 10^4 cells/well. The plate was incubated for 5 h until images were captured by fluorescent microscope (Olympus).

### *In vitro* BBB model

hCMEC/D3 cells and differentiated endothelium derived from iPSCs as previously reported [20] were seeded on the upper chamber of 0.4 μm pore size transwell inserts (#3470, Corning). For non-contact co-culture with HBVP, HBVP were pre-seeded on the underside of the membrane of the inserts.

### TEER and NaF assay

Transendothelial/epithelial electrical resistance (TEER) and the flux of sodium fluorescein (Na-F) were measured to assess the integrity of the BBB model [21]. TEER measurements were taken using an epithelial voltohmmeter (ERS-2) with STX2 electrodes (Millicell, Merck). Resistance values were recorded three times for each well and averaged. These values were corrected for the background resistance of the cell-free insert and the culture medium. The final TEER values were expressed in ohms per square centimeter (Ω·cm^2), calculated by multiplying the net resistance by the surface area of the insert. 100 μg/mL NaF diluted by fresh medium was added to the upper chambers, and the plate was maintained at 37 °C for 1 h. The medium from the lower chambers was collected in a 96-well plate. Then the fluorescence was detected by a microplate reader with the excitation and emission filters set at 488 and 520 nm, respectively. TTR overexpressing hCMEC/D3 cells and controls were seeded on a 12-well plate at a density of 1 × 10^5^ cells per well for 24 hours. Then, cells were treated with 100 μg/mL NaF in fresh medium at 37 °C for 1 h, trypsinized and washed with PBS buffer for flow cytometry analysis (Gallios, Beckman Coulter).

### Protein structure prediction

The protein associations network of TTR was generated using the STRING database. The sequences of TTR and B2M were copied from the UniProt database. The structure prediction of TTR and B2M was generated using AlphaFold Server and visualized using ChimeraX1.8.

### Co-immunoprecipitation (Co-IP)

Co-IP was conducted following the instructions (#88804, Thermo). In brief, cell lysates of hCMEC/D3 were incubated with anti-B2M and anti-TTR **(Table S2)** overnight at 4°C. The antigen/antibody complex was then incubated with protein A/G magnetic beads for 1 hour at RT. Magnetic beads were washed twice with immunoprecipitation lysis/wash buffer, followed by one wash with pure water. The antigen/antibody complex was eluted and collected for further WB assay.

### Fecal microbiota transplantation

Fecal microbiota transplantation was facilitated after 16S rRNA sequencing as previously introduced [16]. Briefly, WT littermates were randomized into two groups after weaning and gavaged with fecal supernatant (200 μL per mouse) of 12-month-old APP/PS1 mice or themselves as control three times a week.

### Behavioral assessments

Behavioral assessments were conducted as reported [22, 23]. Open field test was performed to assess the motor function. Y-maze test was used to assess the spatial working and short- term memory. Novel object recognition test was applied for the innate exploratory behavior and memory assessment. Morris water maze test was employed for analysis of spatial working memory.

### Statistics

Prism 9.0 (GraphPad Software) was utilized for all statistical analyses. The outcomes are expressed as means ± standard error of the mean (SEM). Statistical significance was assessed by unpaired t-test, one-way ANOVA followed by Tukey’s multiple comparisons test or two- way ANOVA analysis. Significance levels are indicated as follows: \**P* < 0.05; \*\**P* < 0.01; \*\*\**P* < 0.001; \*\*\*\**P* < 0.0001.

## Results

### Gut inflammation induces an dynamic hippocampal neurovascular response

Intestinal inflammation was replicated in 2-month-old wild type mice (2m-WT) using a classic dextran sulfate sodium (DSS) model **(Fig. 1A-C)** [17, 24] as a discovery tool, which led to a compromised gut vascular barrier **(Fig. S1)**. With Lectin as vessel markers, the vulnerability of vascular barriers was evaluated by conducting a well-defined extravasated IgG staining analysis [25–27] in the BCSFB and BBB across different brain regions during intestinal inflammation **(Fig. 1D)**. Consistent with the closure of BCSFB during intestinal inflammation reported previously [17], in the choroid plexus of the lateral ventricle, a transient increase of IgG extravasation in BSCFB was observed one day post-treatment (T1, *P* < 0.05), followed by a considerable decrease three days post-treatment (T3, *P* < 0.05), returning to untreated (T0) level, and stabilized until five days post-treatment (T5, *P* = 0.0670) **(Fig. 1E-F)**. However, no significant differences were observed in the permeability of IgG in the choroid plexus of the fourth ventricle **(Fig. 1G-H)**.

**Figure 1.**
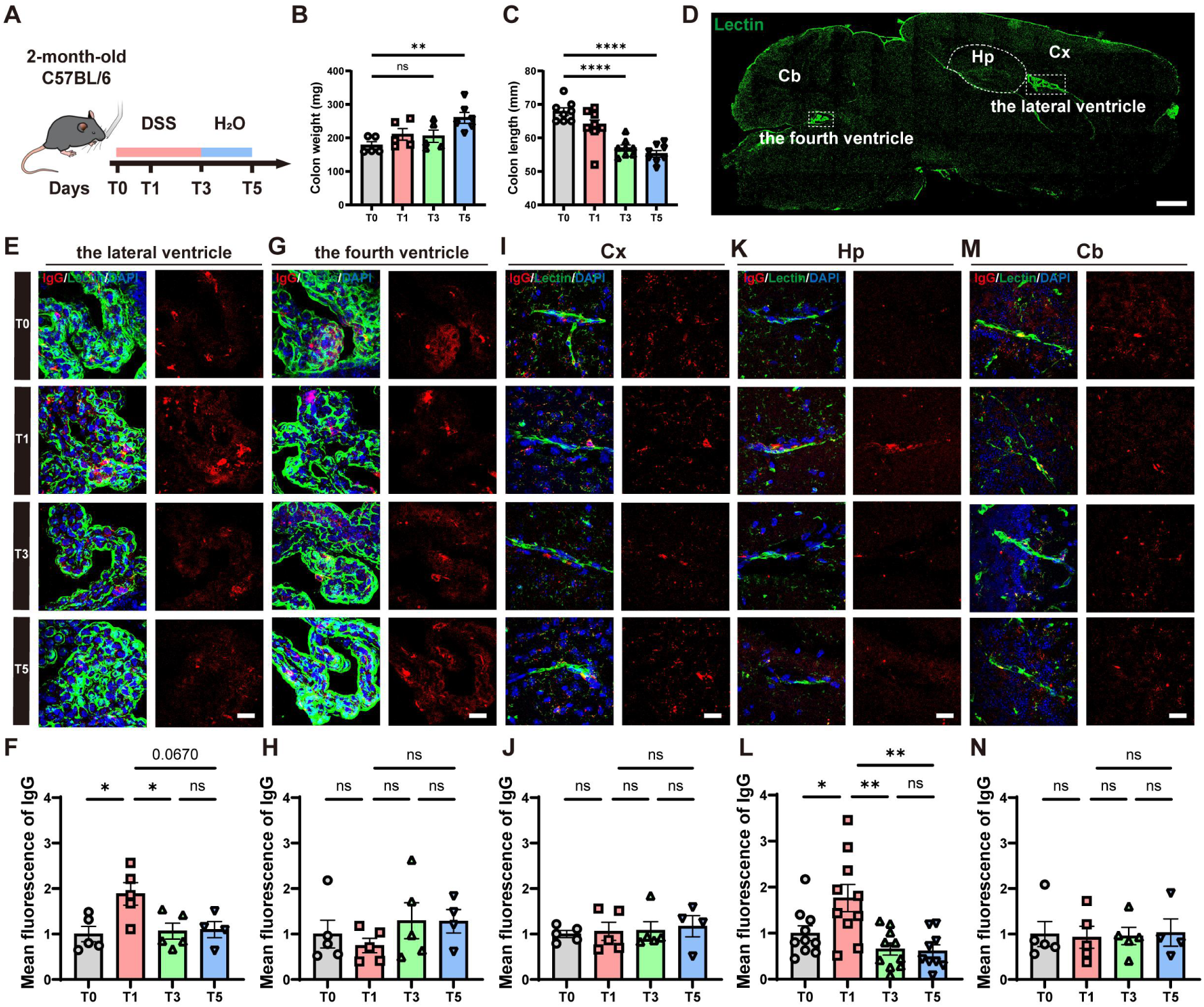
Intestinal inflammation alters the BCSFB and BBB permeability in the lateral ventricle and hippocampus. **(A)** Experimental layout of the intestinal inflammation model induced by DSS administration in drinking water for 1 day (T1) and 3 days (T3), followed by 2 days of regular water (T5) in 2-month-old C57BL/6 mice (2m-WT). **(B)** Colon weight of the mice. N = 5 mice for T0, T1 and T3. N = 6 mice for T5. **(C)** Colon length of the mice. N = 8 mice for each group. **(D)** Representative Z-stack confocal image of blood vessels and choroid plexus labeled by Lectin (green) in the sagittal brain section of mice. Quantitative Z- stack confocal images were taken in the cortex (Cx), hippocampus (Hp), cerebellum (Cb), the lateral ventricle and the fourth ventricle, respectively. Scale bar = 1 mm. **(E-F)** Representative Z-stack confocal images **(E)** and quantification **(F)** of extravasated plasma IgG (red) around choroid plexus labeled by Lectin (green) in the lateral ventricle. N = 5 mice for T0, T1 and T3. N = 4 mice for T5. **(G-H)** Representative Z-stack confocal images **(G)** and quantification **(H)** of extravasated plasma IgG (red) around choroid plexus labeled by Lectin (green) in the fourth ventricle. N = 5 mice for T0, T1 and T3. N = 4 mice for T5. **(I-J)** Representative Z-stack confocal images **(I)** and quantification **(J)** of extravasated plasma IgG (red) around capillaries labeled by Lectin (green) in the Cx. N = 5 mice for T0, T1 and T3. N = 4 mice for T5. **(K-L)** Representative Z-stack confocal images **(K)** and quantification **(L)** of extravasated plasma IgG (red) around capillaries labeled by Lectin (green) in the Hp. N = 10 mice for T0 and T1. N = 11 mice for T3. N = 9 mice for T5. Data from two independent experiments are represented using bar graphs with means ± SEM. **(M-N)** Representative Z- stack confocal images **(M)** and quantification **(N)** of extravasated plasma IgG (red) around capillaries labeled by Lectin (green) in the Cb. N = 5 mice for T0, T1 and T3. N = 4 mice for T5. Comparisons between groups were performed using one-way ANOVA followed by Tukey’s multiple comparisons test. Nuclei were stained with DAPI (blue). Scale bars = 20 µm. ns, no significance, \**P* < 0.05, \*\**P* < 0.01, \*\*\*\**P* < 0.0001.

When evaluating BBB permeability, no significant differences in IgG extravasation were observed over time in the cortex or cerebellum **(Fig. 1I-J, M-N)**. However, in the hippocampus, the extravasated IgG significantly increased at T1 (*P* < 0.05), followed by a significant decline at T3 (*P* < 0.01) **(Fig. 1K-L)**, suggesting a dynamic change exclusively in hippocampal BBB permeability, rather than in that of cortex or cerebellum, induced by modest intestinal inflammation.

In an earlier investigation, we observed an age-dependent BBB disruption associated with enhancing hippocampal transmission of gut-derived amyloid beta (Aβ) via blood [16]. To investigate whether intestinal inflammation also induces hippocampal BBB dynamic regulation in an age-dependent manner, we induced an intestinal inflammation model with DSS in 10m-WT **(Fig. S2A-B)**. In contrast to 2m-WT mice, DSS treatment did not further increase hippocampal IgG extravasation in 10m-WT mice **(Fig. S2C-D)**. Importantly, this does not necessarily indicate a lack of BBB disruption in aged animals, but rather suggests a loss of dynamic responsiveness, potentially due to a higher baseline level of BBB permeability and/or a reduced capacity for further modulation.

Overall, these results suggest that intestinal inflammation transiently, as well as region- specifically regulates not only BCSFB permeability in the choroid plexus of the lateral ventricle, as reported by others, but also BBB permeability in the hippocampus, especially in young mice.

### TTR is induced during the dynamic neurovascular response

To explore the mechanisms underlying the transient shift in hippocampal BBB permeability and elucidate the molecular features of individual cell types in the hippocampus during intestinal inflammation, we performed droplet-based single-cell RNA sequencing (scRNA- seq) without nuclear isolation, focusing on capillary cells rather than neurons, using pooled bilateral hippocampi from three mice per time point as a single sample, with each group replicated twice **(Fig. 2A)**. After quanlity control and doublet removal, we profiled a total of 67,422 hippocampal cells **(Fig. S3 A-C)**. Uniform manifold approximation and projection (UMAP) analysis classified typical CNS cells, specifically neuron, astrocyte, oligodendrocyte progenitor cell (OPC), microglia, along with mesenchymal cells, including endothelium, pericyte, choroid plexus, and smooth muscle cell (SMC) **(Fig. 2B, Fig. S3 D-F)**.

**Figure 2.**
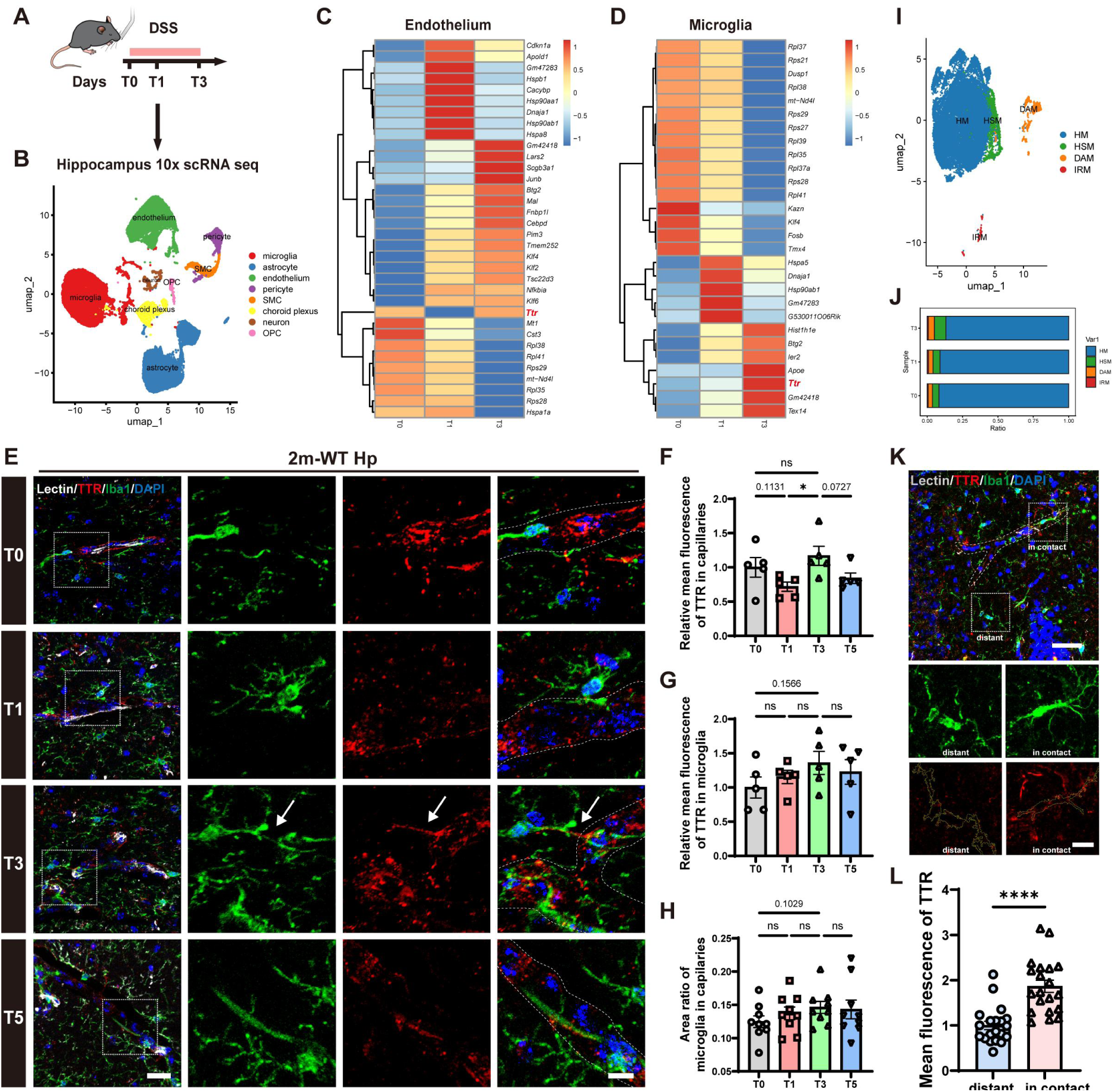
TTR and HSM were remodeled in hippocampal capillaries during intestinal inflammation. **(A)** Hippocampi from mice at T0, T1 and T3 were analyzed by 10x single- cell RNA sequencing (scRNA seq). Bilateral hippocampi of three mice were pooled together as one independent sample, and two repetitive samples were profiled for each time point. **(B)** Visualization of eight major cell types (including microglia, astrocyte, endothelium, pericyte, smooth muscle cell, choroid plexus, neuron, and oligodendrocyte progenitor cell) using UMAP. **(C-D)** Heatmap showing the differentially expressed genes (DEGs) between T0, T1 and T3 in endothelium **(C)** and microglia **(D)**. **(E)** Representative Z-stack confocal images of TTR (red) and microglia labeled by Iba1 (green) around hippocampal capillaries labeled by Lectin (white) of 2m-WT. The merged images in the fourth column display signals in the absence of Lectin. Scale bar = 50 µm (left) and 20 µm (right). **(F)** Quantification of TTR level in the hippocampal capillaries. N = 5 mice for each group. **(G)** Quantification of TTR level in microglia around hippocampal capillaries. N = 5 mice for each group. **(H)** Quantification of the area ratio of microglia in the hippocampal capillaries. The area ratio refers to the proportion of capillary area occupied by microglia, calculated as the total microglial signal within the boundaries of hippocampal capillaries relative to the capillary area. N = 9 mice for each group. **(I)** Visualization of HM, DAM, IRM and HSM using UMAP. **(J)** Cell ratio of the four microglial subtypes at T0, T1 and T3. **(K-L)** Representative Z-stack confocal images **(K)** and quantification **(L)** of TTR (red) in Iba1-positive microglia (green), categorized as capillary-contacting or non-contacting based on Lectin-labeled vasculature (grey) in the Hp. N = 20 mice for each group. Yellow ROIs indicate the microglial shapes. Scale bar = 50 µm (up) and 20 µm (down). Nuclei were stained with DAPI (blue). Comparisons between groups were performed using one-way ANOVA followed by Tukey’s multiple comparisons test for **(F-H)** and unpaired t test for **(L)**. ns, no significance, \**P* < 0.05, \*\*\*\**P* < 0.0001.

Pairwise differential expression analysis across different time points revealed dynamic gene expression changes in individual cell types. In endothelium, *Ttr* notably showed lower level at T1 and returned to baseline at T3 **(Fig. 2C)**, a pattern opposite to hippocampal BBB permeability. A similar upward trend in *Ttr* expression was also observed in microglia **(Fig. 2D)**, pericytes **(Fig. S4A)** and neurons **(Fig. S4C)**, but not in astrocytes **(Fig. S4B)**, suggesting a cell-type-specific regulation of *Ttr* in intestinal inflammation.

Given the neuroprotective roles of TTR [28, 29], we next quantified TTR protein levels in other independent hippocampal tissues of treated 2m-WT during intestinal inflammation. WB results showed that the hippocampal TTR expression significantly increased from T1 to T3 (*P* < 0.05) **(Fig. S4D-E)**, supporting the overall *Ttr* upregulation observed in our hippocampal scRNA-seq data. Although TTR is predominantly synthesized in the choroid plexus in the brain [30], as confirmed by our scRNA-seq data, we also detected lower-level TTR expression across multiple hippocampal cell types. Importantly, this signal contrasts with other choroid plexus-specific marker genes, which remain confined to the choroid plexus **(Fig. 2B, Fig. S4F)**, suggesting that the observed hippocampal TTR expression reflects genuine transcription rather than ambient RNA contamination or sequencing artefacts. To assess TTR protein expression in endothelial cells and microglia, immunofluorescence staining for TTR and Iba1 was performed in human hippocampal sections containing the choroid plexus, revealing detectable TTR signals in hippocampal capillaries and microglia **(Fig. S4G-I)**.

Then, we performed immunofluorescence staining for TTR, capillaries (Lectin) and microglia (Iba1) in brain sections of DSS-treated 2m-WT. We observed that TTR expression in hippocampal capillaries showed a decreased trend at T1 (*P* = 0.1131) and significantly increased at T3 (*P* < 0.05) **(Fig. 2E-F)**. Meanwhile, TTR expression in microglia showed an upward trend, increasing from T0 to T3 (*P* = 0.1566) **(Fig. 2E, G)**, suggesting that TTR in hippocampal capillaries and microglia might be involved in the regulation of BBB reactivity.

Given the age-dependent changes in BBB permeability during intestinal inflammation, we also performed immunofluorescence staining of TTR, capillaries and microglia in DSS- treated 10m-WT. No increase of TTR expression was observed; instead, a declining trend was noted at T3 in both hippocampal capillaries (*P* = 0.2464) and microglia (*P* = 0.0932) **(Fig. S5A-C)**. These results reveal a different TTR variation mode in aged mice, which is consistent with the reduced regulation of hippocampal BBB permeability **(Fig. S2C-D)**, indicating that the function of TTR is likely to decline with aging.

### A vascular-associated TTR-high microglial state accompanies neurovascular adaptation

To further explore the potential roles of microglia in BBB reactivity, subcluster analysis of microglia was performed, identifying homeostatic microglia (HM) [31, 32], disease- associated microglia (DAM) [33–35], interferon-responsive microglia (IRM) [36], and a novel subcluster with higher *Ttr* expression **(subcluster 3; Fig. S6A-D)**. Given the association between dynamic *Ttr* expression changes and intestinal inflammation, we designated this microglial state as “high-sensitive microglia” (HSM) **(Fig. 2I)**. Heatmap highlighted the top 7 differential expressed genes (DEGs) among the four states, with HSM exhibiting DEGs associated with vascular regulation [37, 38], neuroprotection[39, 40], and energy metabolism [41] **(Fig. S6E)**. GO enrichment analysis further validated the classification of HM, DAM, and IRM **(Fig. S6F)**. Besides, 6 DEGs in HSM were overlapped with vascular gene sets **(Fig. S6G, Table S3)**, further supporting its potential vascular association.

Notably, the proportion of HSM showed an increasing trend during intestinal inflammation **(from 0.045 at T0 to 0.081 at T3; Fig. 2J)**. To correlate the HSM subtype with vascular- associated microglia, microglia touching or wrapping around hippocampal capillaries were further analyzed. The area ratio of microglia in contact with vasculature increased from T0 to T3 in treated 2m-WT (*P* = 0.1029), coinciding with the increased proportion of HSM, but not in 10m-WT **(Fig. 2E, H and Fig. S5A, D)**. Comparative analysis of TTR protein levels across two microglial subpopulations — those in contact with the vasculature versus those distant from it—further confirmed that microglia adjacent to capillaries exhibit significantly higher TTR expression. (*P* < 0.0001, **Fig. 2K-L)**.

These results reveal a potential vascular-associated microglial state, marked by higher TTR expression, indicating a possible association with vascular sensing during acute intestinal inflammation.

### Overexpression of hippocampal TTR rescues the BBB leak induced by LPS-mediated GBA dysfunction

To verify the protective role played by hippocampal TTR during GBA dysfunction, pAAV5- CBh-mTtr-EGFP and pAAV5-CBh-NC-EGFP were stereotaxically injected into the hippocampus of 2m-WT (abbreviated as mTtr and NC group) **(Fig. 3A)**. Given the broad cellular tropism of AAV5 in the central nervous system, multiple hippocampal cell types were transduced following injection. In the present study, we focused our analysis on vascular endothelial cells and HSM, based on our prior findings. After 6 weeks of injection, we observed an increased TTR in capillaries (*P* < 0.01) and HSM (*P* < 0.01) **(Fig. 3B-E)**. Consistent with **Fig. 2H**, we observed a significantly increased area ratio of HSM in hippocampal capillaries (*P* < 0.05) **(Fig. 3C, F)**.

**Figure 3.**
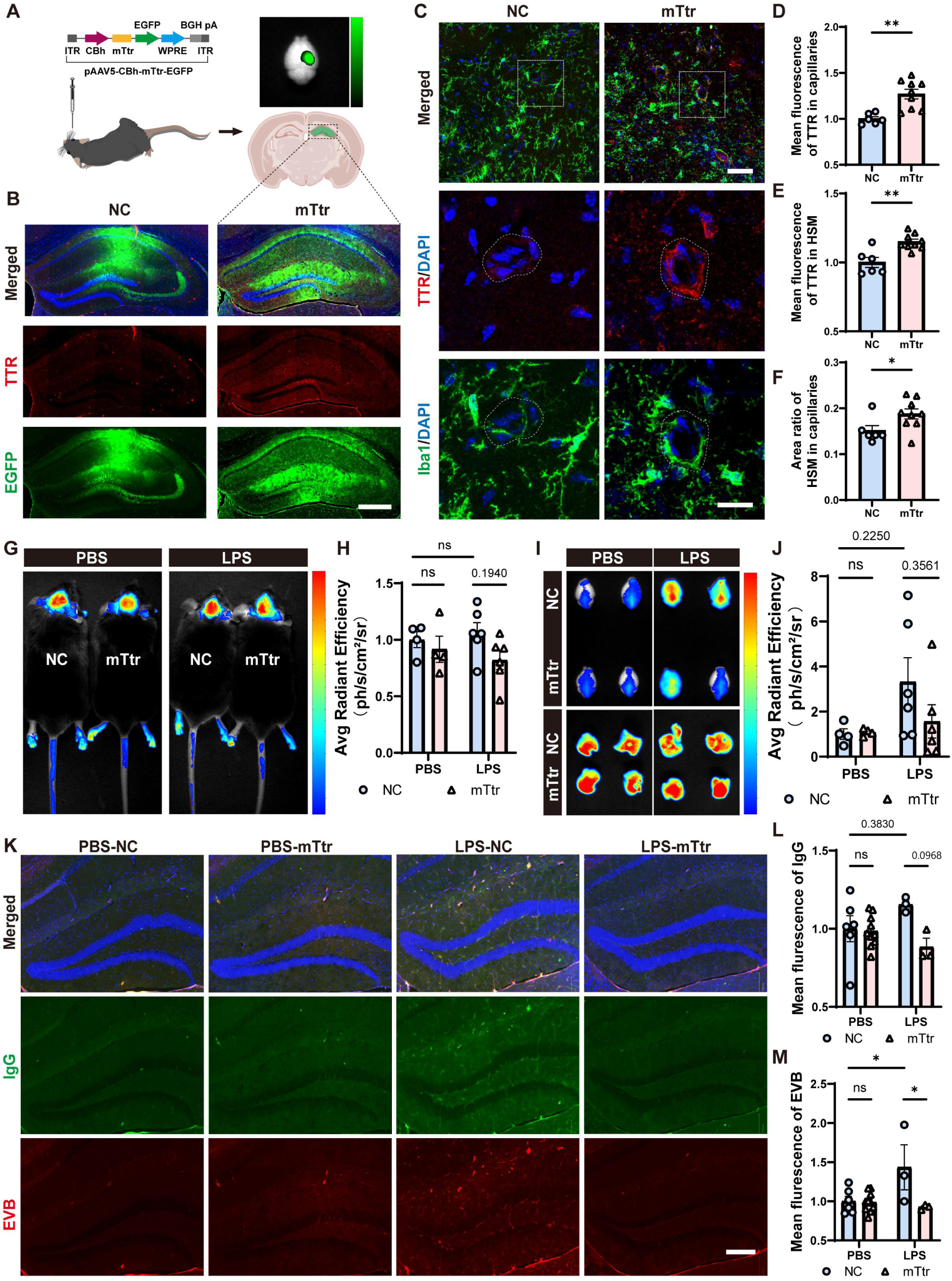
Overexpression of TTR in hippocampus alleviated LPS-induced BBB leakiness. **(A)** pAAV5-CBh-mTtr-EGFP (mTtr) and pAAV5-CBh-NC-EGFP (NC) were stereotaxically injected into the right Hp of 2m-WT mice. **(B)** Representative IF images of TTR (red) and EGFP (green) in Hp 6 weeks after the injection. Scale bar = 500 µm. **(C)** Representative Z- stack confocal images of TTR (red) and microglia labeled by Iba1 (green) around the hippocampal capillaries in the mice injected with NC and mTtr. Scale bar = 50 µm (above) and 20 µm (below). (D-E) Quantification of TTR in hippocampal capillaries **(D)** and HSM **(E)** of NC and mTtr. N = 6 mice for NC and N = 9 mice for mTtr. **(F)** Quantification of the area ratio of HSM in hippocampal capillaries. N = 6 mice for NC and N = 9 mice for mTtr. **(G-H)** Representative *in vivo* images **(G)** and quantification **(H)** of intravenously injected EVB in the brain of NC and mTtr after the intraperitoneal injection of PBS or LPS. N = 4 mice for NC and mTtr in the PBS group and N = 6 mice for NC and mTtr in the LPS group. **(I-J)** Representative ex vivo images of the extravasated EVB in the brain and liver **(I)** and quantification **(J)** of the ratio of extravasated EVB in the brain related to that in liver. N = 4 mice for NC and mTtr in the PBS group and N = 6 mice for NC and mTtr in the LPS group. **(K-M)** Representative IF images **(K)** and quantifications of extravasated plasma IgG (green) **(L)** and EVB (red) **(M)** around the hippocampal capillaries. N = 6 mice for NC and N = 10 mice for mTtr in the PBS group. N = 3 mice for NC and mTtr in the LPS group. Scale bar = 200 µm. Only the right hippocampi were analyzed in **(B-F)** and **(K-M)**. Nuclei were stained with DAPI (blue). Comparisons between groups were performed with two-way Anova for multiple groups and unpaired t test for two groups. ns, no significance, \**P* < 0.05, \*\**P* < 0.01.

Intraperitoneal LPS injection was selected for subsequent in vivo functional studies of TTR because it induces a rapid, synchronized, and systemic inflammatory response, enabling precise temporal assessment of BBB changes. The permeability of hippocampal BBB was evaluated by EVB as previously reported [42–44] following intraperitoneal injection of PBS or LPS. Although no significant difference of *in vivo* signal was observed in the brain between NC and mTtr mice after PBS injection, after LPS, overexpression of TTR, to some extent, alleviated LPS-induced BBB leakiness (*P* = 0.1940) **(Fig. 3G-H)**. After perfusion, the same trend was observed when extravasated EVB fluorescence in the brain was quantified relative to that in the liver (*P* = 0.2250 between NC injected with PBS or LPS, *P* = 0.3561 between LPS-treated NC and mTtr) **(Fig. 3I-J)**. Similarly, quantifications of fluorescence of IgG and EVB across all the hippocampal subregions further supported the optical imaging findings, with a decreasing trend of IgG (0.0968) and significantly decreased EVB (*P* < 0.05) observed in the overexpression mice in LPS group (*P* < 0.05) **(Fig. 3K-M)**.

These results suggest that overexpression of TTR in the hippocampus ameliorates the hippocampal BBB leakage during GBA dysfunction.

### Downregulation of TTR in the hippocampus increases BBB permeability

To reversely validate our investigation in **Fig. 3**, we further explored whether reduced TTR in the hippocampus directly leads to BBB leakiness. For this purpose, pAAV5-U6- mTtr[shRNA]-mCherry was stereotaxically injected into the hippocampus of 2m-WT (shRNA) and pAAV5-U6-scramble-mCherry was used as negative control (NC) **(Fig. 4A)**. After 6 weeks, TTR expression was downregulated in the hippocampus, especially in capillaries and HSM (*P* < 0.05) **(Fig. 4B-E)** in the shRNA group. Remarkably, the area ratio of HSM significantly decreased (*P* < 0.01) **(Fig. 4C, F)**.

**Figure 4.**
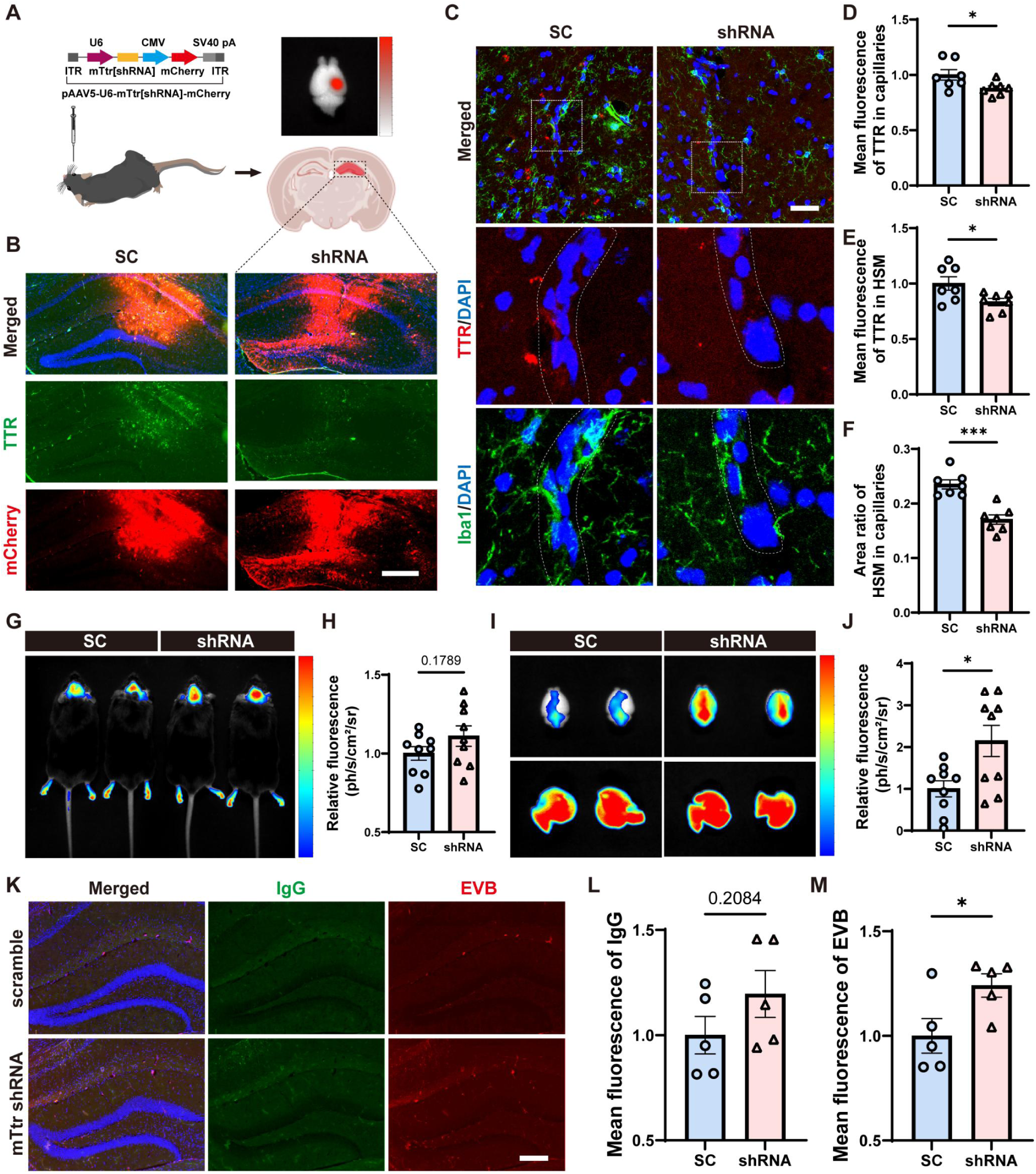
Downregulation of TTR in hippocampus increases the permeability of BBB. **(A)** pAAV5-U6-mTtr[shRNA]-mCherry (shRNA) and pAAV5-U6-scramble-mCherry (SC) were stereotaxically injected into the right Hp of 2m-WT. **(B)** Representative IF images of TTR (green) and mCherry (red) in Hp 6 weeks after the injection. Scale bar = 500 µm. **(C)** Representative Z-stack confocal images of TTR (red) and microglia labeled by Iba1 (green) around the hippocampal capillaries in the mice injected with SC and shRNA. Scale bar = 50 µm (above) and 20 µm (below). **(D-E)** Quantification of TTR in hippocampal capillaries **(D)** and HSM **(E)** of SC and shRNA. N = 7 mice for each group. **(F)** Quantification of the area ratio of HSM in hippocampal capillaries of SC and shRNA. N = 7 mice for each group. **(G-H)** Representative in vivo images **(G)** and quantification **(H)** of intravenously injected EVB in the brain of SC and shRNA. N = 9 mice for each group. **(I-J)** Representative *ex vivo* images of the extravasated EVB in the brain and liver **(I)**, and quantification of extravasated EVB in the brain related to that in the liver **(J)**. N = 9 mice for each group. **(K-M)** Representative IF images **(K)** and quantifications of extravasated plasma IgG (green) **(L)** and EVB (red) **(M)** around the hippocampal capillaries. N = 5 mice for each group. Scale bar = 200 µm. Only the right hippocampi were analyzed in **(B-F)** and **(K-M)**. Nuclei were stained with DAPI (blue). Comparisons between groups were performed with unpaired t test. ns, no significance, \**P* < 0.05, \*\*\**P* < 0.001.

The permeability of hippocampal BBB was evaluated by EVB. *In vivo* imaging showed an increasing trend of EVB (*P* = 0.1789) in the brain of shRNA **(Fig. 4G-H)**, indicating increased permeability of BBB. After perfusion, *ex vivo* imaging further validated this observation, with a significant increase of the extravasated EVB in the brain/liver (*P* < 0.05) **(Fig. 4I-J)**. Consistently, more IgG (*P* = 0.2084) and EVB (*P* < 0.05) were detected across the entire hippocampal region of shRNA **(Fig. 4K-M)**, indicating that downregulation of TTR in the hippocampus increases BBB permeability.

### Overexpression of TTR drives endothelial migration and angiogenesis

To further explore the mechanisms by which TTR and HSM promote vascular remodeling, we examined intercellular ligand-receptor interactions among extracted HM, DAM, IRM, HSM and endothelium **(Fig. S7A)** using CellChat analysis. The complex interacting network was visualized in a circular plot **(Fig. S7B)**. The bubble plot illustrated various interacting gene pairs, showing that the *Ptn-Ncl* interaction was exclusively observed during the interaction from HSM to endothelium (highlighted in red) **(Fig. S7C)**.

PTN has been reported to be involved in endothelial migration and angiogenesis, mediated by NCL in endothelium [45, 46]. In response to certain physiological or pathological stimuli, controlled angiogenesis supports vascular restoration and functional recovery [47–50]. To test this potential pathway induced by TTR, human TTR (hTTR) overexpression model was separately constructed in human brain endothelium cell line (hCMEC/D3) and human microglia cell line (HMC3) by lentiviral infection **(Fig. S8A-C)**. Wound healing assay and tube formation assay were conducted to assess endothelial migration and *in vitro* angiogenesis. The results demonstrated that both direct overexpression of hTTR in hCMEC/D3 (**Fig. 5A-E**) and exposure to conditioned medium from hTTR-overexpressing HMC3 (**Fig. 5F-J**) significantly enhanced migration and tube formation in hCMEC/D3. These findings are consistent with previous reports describing the pro-angiogenic properties of TTR [51, 52] and further suggest that TTR, potentially in concert with HSM, may contribute to the maintenance of vascular function.

**Figure 5.**
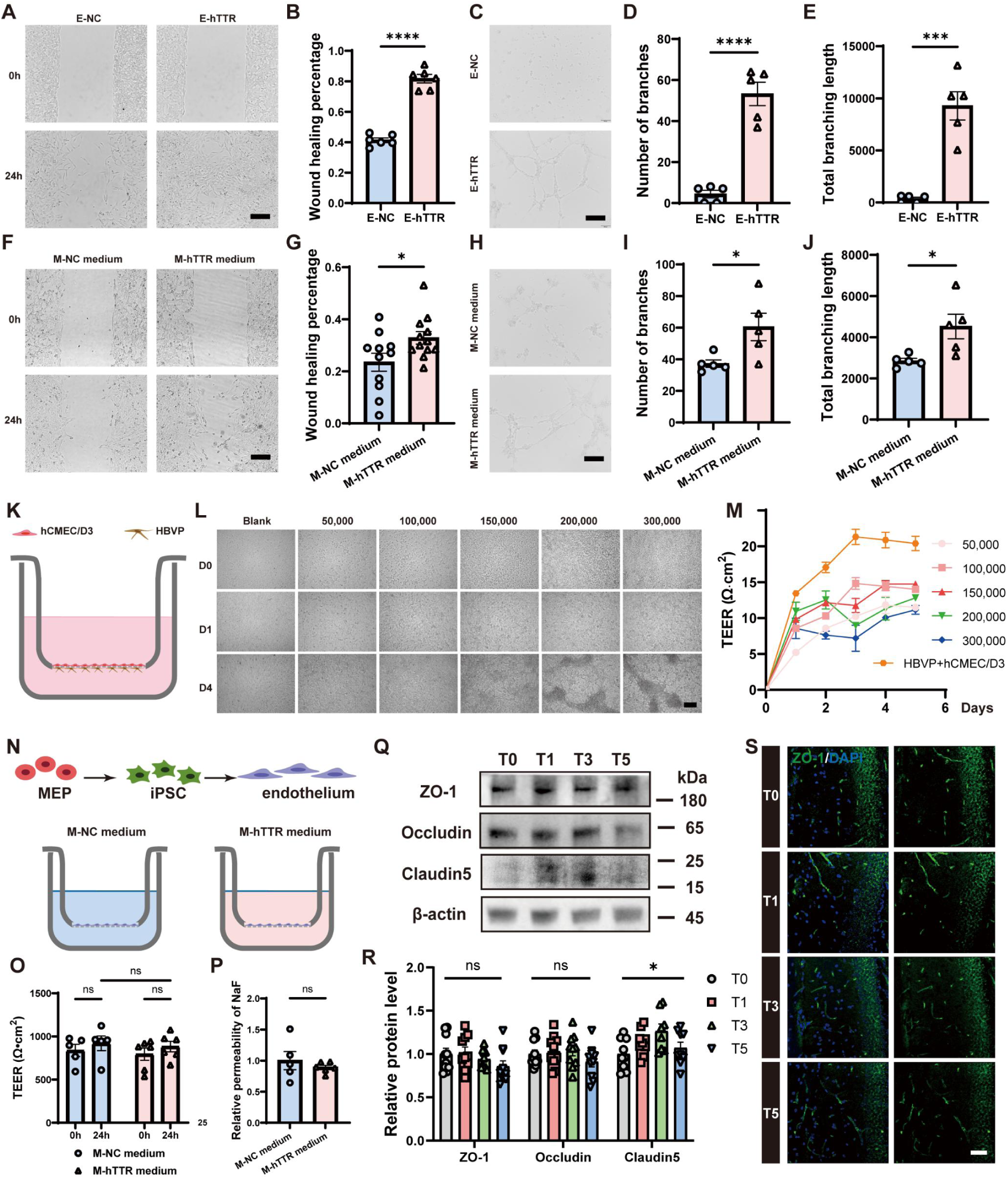
Overexpression of TTR in cell lines promotes wound healing and tube formation, and BBB *in vitro* model. **(A-B)** Representative wound healing images **(A)** and quantification of wound healing percentage at 24h **(B)** in hCMEC/D3 infected with hTTR expressing lentivirus (E-hTTR) and negative lentivirus vectors (E-NC). Scale bars = 200 µm. N = 6 wells. **(C-E)** Representative tube formation images **(C)** and quantifications of the number of branches **(D)** and total branching length **(E)** in E-NC and E-hTTR. Scale bars = 200 µm. N = 5 wells. **(F-G)** Representative wound healing images **(F)** and quantification of wound healing percentage at 24h **(G)** in hCMEC/D3 cultured with conditional medium from hTTR-overexpressing HMC3 (M-hTTR medium) and controlled HMC3 (M-NC medium). Scale bars = 200 µm. N = 12 wells. **(H-J)** Representative tube formation images **(H)** and quantifications of the number of branches **(I)** and total branching length **(J)** in M-NC medium and M-hTTR medium groups. Scale bars = 200 µm. N = 5 wells. **(K)** Schematic diagram of BBB model in transwell using hCMEC/D3 and HBVP. **(L)** Representative images of different numbers of hCMEC/D3 cells cultured on 0.4 μm PET membrane from Day 0 (D0) to Day 4 (D4). Scale bar = 200 µm. **(M)** Quantification of TEER in transwell cultured with different numbers of hCMEC/D3 cells and non-contact co-culture of HBVP and hCMEC/D3 from D0-D5. **(N)** Schematic diagram of BBB model in transwell using endothelium induced from induced pluripotent stem cells (iPSC) derived from megakaryocyte-erythroid progenitor (MEP). **(O)** Quantification of TEER in transwell cultured with induced human endothelium before (0h) and 24h after exposure to medium from HMC3 infected with hTTR expressing lentivirus (M-hTTR medium) and medium from HMC3 infected with negative lentivirus vectors (M-NC medium). N = 5 wells for M-NC medium and N = 6 wells for M-hTTR medium. **(P)** Quantification of permeability of NaF at 1h in M-NC medium and M-hTTR medium. N = 5 wells for M-NC medium and N = 6 wells for M-hTTR medium. **(Q-R)** Representative WB images **(Q)** and quantifications **(R)** of the expressions of ZO-1, Occludin and Claudin5 in Hp. For ZO-1 and occludin, N = 10 mice for each group. For Claudin5, N = 9 mice for each group. **(S)** Representative Z-stack confocal images of ZO-1 (green) in the Hp to support the WB quantification. Nuclei were stained with DAPI (blue). Scale bars = 50 µm. Comparisons between groups were performed with ordinary one-way ANOVA test for **(R)** and unpaired t test for others. ns, no significance, \**P* < 0.05, \*\*\**P* < 0.001, \*\*\*\**P* < 0.0001.

We further established an *in vitro* BBB model to investigate whether TTR affects tight junctions. Either varying the seeding density of hCMEC/D3 cells alone or co-culturing them in a non-contact setup with human brain pericytes (HBVP) resulted in relatively low TEER values in the Transwell assay **(Fig. 5K-M)**. We previously induced vascular endothelial cells generated from human induced pluripotent stem cells (iPSC) derived from megakaryocyte- erythroid progenitor [53]. Thus, the vascular endothelial cells were used instead to establish BBB *in vitro* **(Fig. 5N)**. After culture with the conditioned media from hTTR-overexpressing HMC3 or controlled HMC3, no significant differences were observed in the TEER or NaF permeability in iPSC-differentiated BBB model **(Fig. 5O-P)**, suggesting that while HSM may be associated with angiogenic processes, its involvement in the regulation of tight junctions appears less evident. To validate this observation *in vivo*, three tight junction proteins were quantified in the hippocampus of treated 2m-WT. While Claudin 5 dynamically varied (*P* < 0.05), ZO-1 and Occludin showed no considerable shifts during the intestinal inflammation **(Fig. 5Q-S)**. Nevertheless, the alteration mode of the hippocampal Claudin 5 did not correspond to the IgG extravasation. Collectively, these results suggest that the intercellular tight junctions are not the primary contributors to the observed dynamic alterations in hippocampal BBB permeability.

### TTR associates with beta2-microglobulin (B2M), potentially contributing to reduced neonatal Fc receptor (FcRn)-mediated endocytosis

Given that paracellular pathways do not appear to account for the changes in BBB permeability, we next hypothesized TTR modulates endothelial transcellular transport. To specifically explore the mechanisms underlying the potential TTR-mediated regulation of endothelial transcytosis, we first analyzed the protein-protein associations network of TTR from the STRING database **(Fig. 6A)**, and focused on B2M, which binds with FCGRT as a subunit of FcRn [54] **(Fig. 6B)**. FcRn mediates transcytosis of both albumin and IgG in capillary endothelium [55]. According to the AlphaFold Server, TTR was predicted to bind with B2M by several H-bonds. Lys6, Tyr10, Asp53, and Ser55 were shared binding sites in B2M **(Fig. 6B)**, indicating competitive binding between TTR and FCGRT. Experimentally, Co-IP of TTR and B2M in hCMEC/D3 lysates was performed to validate that TTR and B2M associate with each other **(Fig. 6C)**. Colocalization of these two proteins was also observed on the membrane of endothelial cells **(Fig. 6D)**. Additionally, overexpression of hTTR in hCMEC/D3 significantly decreased the colocalization of B2M and FCGRT (*P* < 0.05) **(Fig. 6E-F)**. Finally, overexpression of hTTR significantly decreased the endocytosis of NaF in hCMEC/D3 (*P* < 0.05) **(Fig. 6-H)**, suggesting that TTR likely reduces endothelial transcytosis via FcRn receptors.

**Figure 6.**
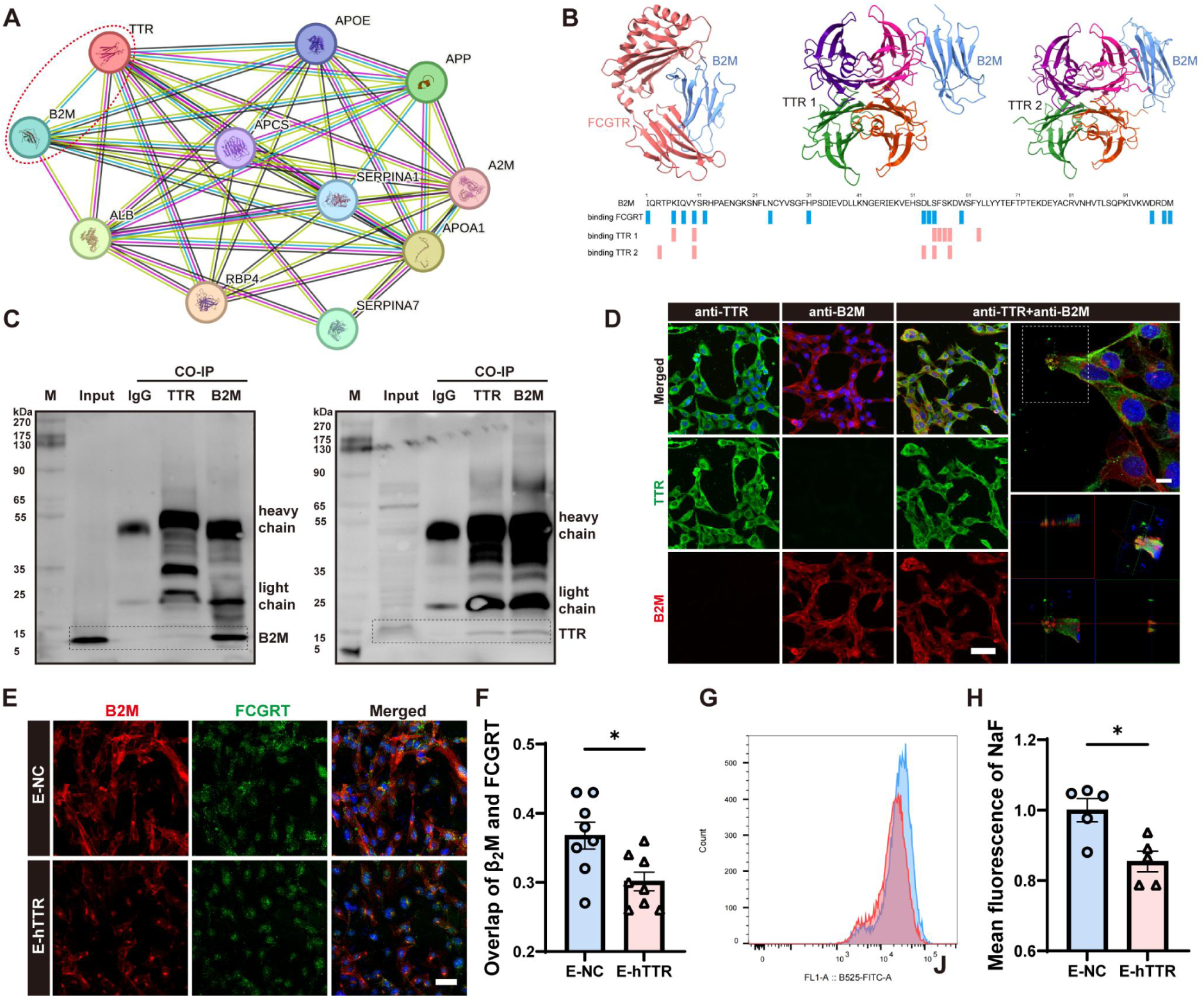
TTR binds with beta2-microglobulin (B2M) and reduces neonatal Fc receptor (FcRn)-mediated endocytosis. **(A)** Protein-protein associations network of TTR from the STRING database. **(B)** 3D Visualization of FcRn composed of FCGRT and B2M and TTR binding to B2M. Sequence annotation showed the binding sites of FCGRT (blue) and TTR (pink) in B2M, respectively. The interaction data of TTR and B2M was generated twice by AlphaFold Server and visualized by ChimeraX. **(C)** WB images of B2M (left) and TTR (right) in Input (hCMEC/D3 lysate) and CO-IP samples using isotype control antibody (IgG) and antibodies against B2M and TTR. The heavy chain of antibodies is ≈ 55kDa and the light chain of antibodies is ≈ 25kDa. **(D)** Representative Z-stack confocal images of colocalization of TTR (green) and B2M (red) in hCMEC/D3. Scale bar = 50 µm (left) and 10 µm (right). **(E-F)** Representative Z-stack confocal images **(E)** of B2M (red) and FCGRT (green) and quantification **(F)** of colocalization of these two proteins in hCMEC/D3. Scale bar = 50 µm. N = 8 wells. **(G-H)** Representative flow cytometry histogram plot **(G)**, and quantification **(H)** showing mean fluorescence of NaF in E-NC and E-hTTR. N = 5 wells. Nuclei were stained with DAPI (blue). Comparisons between groups were performed with unpaired t test. \**P* < 0.05.

### Decreased hippocampal TTR in AD model mice and AD patients

Previous research suggested increased permeability of BBB with aging [56–58] and AD [59, 60]. To investigate whether TTR is involved in BBB dysfunction associated with aging and AD, we performed immunofluorescence staining for microglia and TTR in the hippocampus of 2m-WT, 12m-WT and 12-month-old APP/PS1 mice (12m-APP/PS1). In WT mice, a decline in hippocampal capillary TTR (*P* = 0.1166) was observed with aging, which may account for the impaired dynamic regulation of hippocampal BBB permeability during acute inflammation. In 12m-APP/PS1, a further decrease in TTR level was observed in hippocampal capillaries (*P* < 0.05) and HSM (*P* = 0.0530). However, no significant differences in the area ratio of HSM in hippocampal capillaries were observed among the three groups **(Fig. 7A-D)**.

**Figure 7.**
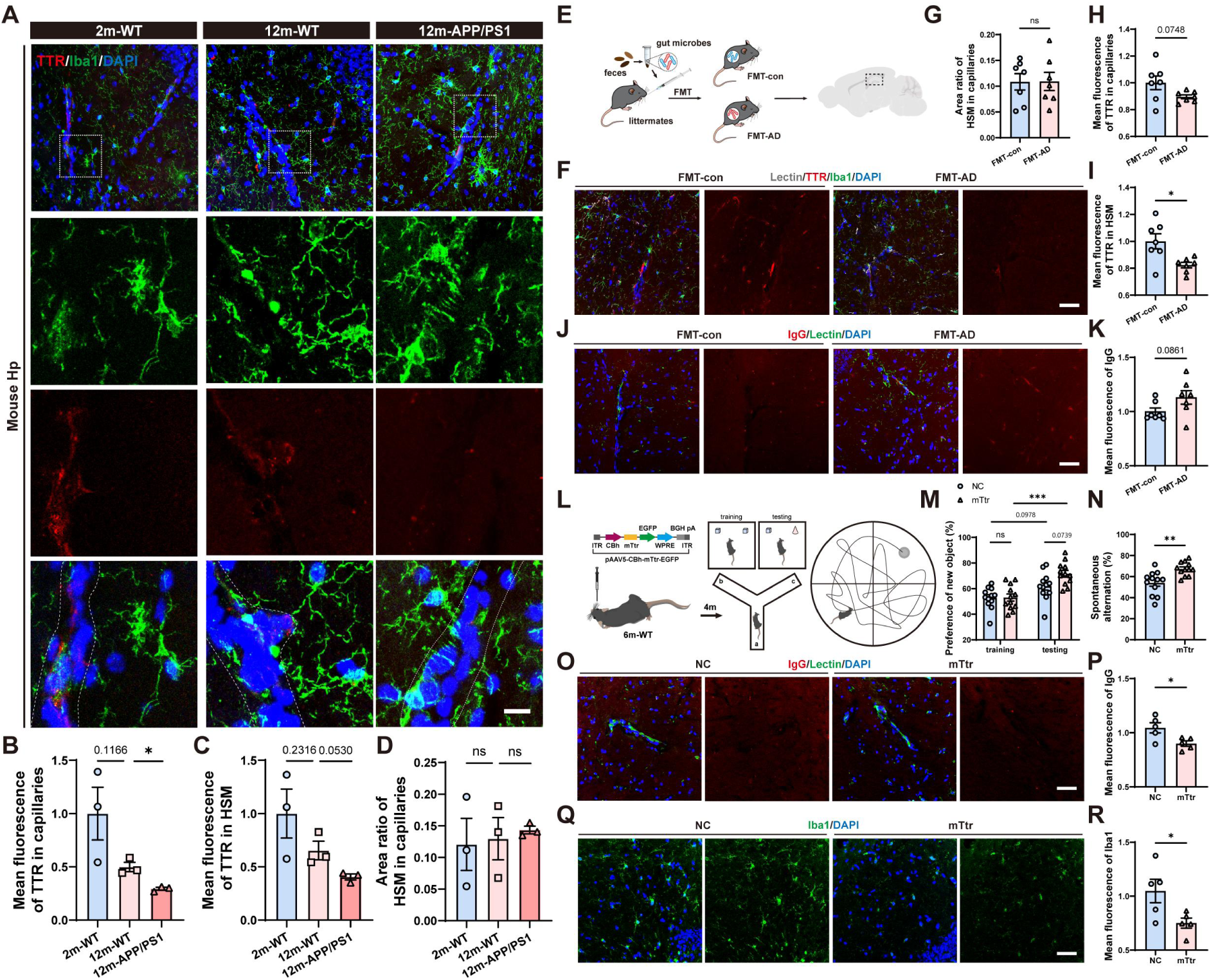
Downregulation of TTR in AD model mice and the therapeutic potential in cognitive impairment, BBB dysfunction and neuroinflammation by restoration of TTR. **(A)** Representative Z-stack confocal images of TTR (red) and microglia labeled by Iba1 (green) around mouse hippocampal capillaries. **(B-C)** Quantification of TTR in hippocampal capillaries **(B)** and HSM **(C)** of 2m-WT, 12m-WT and 12m-APP/PS1. N = 3 mice for each group. **(D)** Quantification of the area ratio of HSM in hippocampal capillaries. N = 3 mice for each group. **(E)** Fecal microbiota from 12m-APP/PS1 were gavaged to newly weaned C57BL/6 mice (FMT-AD). Fecal microbiota from newly weaned C57BL/6 mice were gavaged to themselves as control (FMT-con). **(F)** Representative Z-stack confocal images of TTR (red) and microglia labeled by Iba1 (green) around the mouse hippocampal capillaries labeled by Lectin (grey). **(G)** Quantifications of the area ratio of HSM in hippocampal capillaries of FMT-con and FMT-AD. N = 7 mice for each group. **(H-I)** Quantifications of TTR in hippocampal capillaries **(H)** and HSM **(I)** of FMT-con and FMT-AD. N = 7 mice for each group. **(J-K)** Representative Z-stack confocal images **(F)** and quantification **(G)** of extravasated IgG (red) around hippocampal capillaries labeled by Lectin (green). N = 7 mice for each group. **(L)** pAAV5-CBh-mTtr-EGFP (mTtr) and pAAV5-CBh-NC-EGFP (NC) were stereotaxically injected into the right Hp of 6-month-old C57BL/6 mice. After 4 months, mice were conducted Novel object recognition, Y-maze and Morris’ water maze analysis. **(M-N)** Quantification of preference for new object in Novel object recognition **(M)** and spontaneous alteration in Y-maze **(N)**. N = 12 mice for each group. **(O-P)** Representative Z- stack confocal images of capillary labeled by Lectin (green) and IgG (red) **(O)**, and quantification of IgG extravasation **(P)** in Hp of 10-month-old NC and mTtr. N = 5 mice for each group. **(Q-R)** Representative Z-stack confocal images of microglia labeled by Iba1 (green) **(Q)** and quantification of Iba1 expression **(R)** in Hp of 10-month-old NC and mTtr. N = 5 mice for each group. Cell nuclei were stained with DAPI (blue). Scale bar = 50 µm. Comparisons between groups were performed with unpaired t test and two-way Anova for **(M)**. no significance, \**P* < 0.05, \*\**P* < 0.01, \*\*\*\**P* < 0.0001.

A growing body of research also suggests the correlation between the dysbiotic microbiota and AD pathogenesis [61–66]. We previously induced early AD-like cognitive impairment and neuropathology by fecal microbiota transplantation (FMT) in young WT mice [16] **(Fig. 7E)**. To test whether TTR was influenced in FMT-induced early AD pathogenesis, the same analysis was performed. The results demonstrated that FMT decreased TTR expression in hippocampal capillaries (*P* = 0.0748) and HSM (*P* < 0.05) **(Fig. 7F, H-I)**, along with increased leakiness of plasma IgG (*P* = 0.0861) **(Fig. 7J-K),** indicating a potential contribution of gut microbiota to dynamic regulation of hippocampal BBB permeability via TTR.

To investigate the TTR alteration in human hippocampus, we stained TTR and Iba1 in young healthy controls (young), old healthy controls (old), and AD patients. Contrary to the results in WT mice, significantly increased TTR expression was observed in hippocampal capillaries (P < 0.01) and HSM (P < 0.05) with aging in healthy controls **(Fig. S9A-C)**, suggesting a different evolution of TTR with aging in human hippocampal capillaries. On the other hand, aligned with the data in 12m-APP/PS1, in AD patients, hippocampal TTR expression was lower in the capillaries (P = 0.0936) and HSM (P < 0.1172) **(Fig. S9A-C)**, indicating a compromised hippocampal TTR function in AD process.

### Restoring TTR partially improves age-associated BBB dysfunction and cognition

To investigate the potential therapeutic effect of targeting TTR on aging and neurodegeneration, pAAV5-CBh-mTtr-EGFP and pAAV5-CBh-NC-EGFP were injected stereotaxically into the hippocampus of 6m-WT (abbreviated as mTtr and NC). Four months after injection, there were no obvious alterations in weight or motor activity **(Fig. S10A-F)**. Cognitive impairment matrices, including New Object Recognition, Y-maze and Morris water maze **(Fig. 7L)** were assessed. In training stage of New Object Recognition, both NC and mTtr exhibited approximately 50% preference for the two same objects. In contrast, in the testing stage, mTtr showed a significantly higher preference for the new object (*P* = 0.0739) **(Fig. 7M and Fig. S10G)**. Consistently, comparison between the training and testing stages demonstrated an enhanced novel object preference following TTR overexpression (*P* < 0.001), supporting improved recognition memory. Additionally, the spontaneous alteration in mTtr was significantly higher in Y-maze (*P* < 0.01) **(Fig. 7N and Fig. S10H)**. However, there were no significant differences in mean entries into platform (*P* = 0.6946) and mean time in platform zone (*P* = 0.2254) in Morris Water Maze **(Fig. S10I-K)**. When the brains were assessed histologically, significantly decreased extravasated IgG (*P* < 0.05) **(Fig. 7O-P)** and Iba1 expression in microglia (*P* < 0.05) **(Fig. 7Q-R)** were detected in the hippocampus of mTtr. Given that AAV5-mediated transduction is not restricted to a single cell type, neuronal TTR expression may also contribute to these protective effects **(Fig. S10L)**, although the present study primarily focused on vascular-associated mechanisms. Together, these results suggest that targeting declined TTR improves BBB dysfunction, thereby significantly preventing neuroinflammation and cognitive impairments in aged mice.

## Discussion

Loss of BBB integrity during intestinal inflammation has been well documented [67], with gut-derived inflammatory mediators such as LPS recognized as key contributors. However, the precise mechanisms through which gut inflammation influences brain homeostasis via the GBA remain largely unresolved. A recent study revealed a dynamic, transient closure of the vascular barrier in the choroid plexus, BCSFB, as a novel protective response to DSS- induced intestinal inflammation [17], highlighting that barrier regulation itself may serve as an adaptive defense mechanism.

In the present study, we not only confirmed this intriguing phenomenon but also refined its anatomical specificity—demonstrating that such barrier restoration occurs exclusively in the choroid plexus of the lateral ventricle. More importantly, we discovered that this barrier remodeling is not confined to the choroid plexus. A distinct, previously unrecognized protective restoration of the BBB occurs in the hippocampus in response to gut inflammation. Remarkably, the spatiotemporal dynamics of hippocampal BBB regulation parallel those of the BCSFB, suggesting that gut inflammation triggers coordinated yet region-specific vascular responses within the brain. Furthermore, we found that the hippocampal BBB remodeling is age-dependent—implying that the aging brain, particularly regions critical for learning and memory [68], may be disproportionately vulnerable to gut-derived inflammatory insults.

BBB is traditionally thought to be composed of endothelium, with highly regulated tight junctions, pericytes, and astrocytes [69]. Although the involvement of microglia in regulating BBB integrity remains controversial, especially at physiological conditions [70], their indispensable function in immune surveillance and response influences BBB function [71–73], particularly in the context of neuroinflammation and neurodegenerative diseases. The advent of scRNA-seq technology has further advanced the study of microglia, revealing various microglial subtypes [74], as their function may be regulated by multiple factors, including disease type, age and spatial localization. Our findings identified a novel subpopulation of microglia governed by the GBA, which expresses higher TTR, highlighting a previously unrecognized link between microglial heterogeneity and hippocampal BBB changes during intestinal inflammation.

Tight junction proteins, essential for the integrity of BBB, regulate paracellular permeability [75]. Under normal conditions, tight junctions strictly restrict paracellular transport across the BBB, allowing only small ions and solutes to pass through regulated pathways. Instead, most molecules cross via transcellular routes such as receptor-mediated transcytosis, caveolae- mediated endocytosis, and, to a lesser extent, macropinocytosis [76–79]. Our findings on tight junction proteins suggested that modest intestinal inflammation is insufficient to regulate hippocampal BBB tight junctions in a short time, especially in young, healthy mice. Instead, we demonstrated that the interaction of TTR with B2M and associated FcRn-mediated endocytosis, likely contributed to the dynamic regulation of hippocampal permeability during intestinal inflammation. The detailed mechanisms require further investigation and several key questions remain to be fully understood, including 1) whether TTR binds to B2M in its monomeric or homo-tetrameric form, as well as the structure of the TTR-B2M complex and its binding sites, 2) how TTR regulates FcRn subunit assembly and FcRn recycling, and 3) the potential functions of TTR-B2M complex in the blood.

Earlier investigations indicate that TTR is a protein primarily produced in the liver and choroid plexus of the brain. It functions as a transporter of thyroid hormones (thyroxine, T4) and retinol (vitamin A) by binding to them and facilitating their transport in the bloodstream and cerebrospinal fluid [80, 81]. Mutations in *TTR* gene are associated with amyloid deposition, predominantly affecting peripheral nerves or the heart [82]. Recent research has suggested that by binding to Aβ [83] and facilitating the clearance of Aβ from the brain [14], TTR reduces its ability to form plaques, thereby mitigating the neurotoxic effects of Aβ. Several studies have reported reduced TTR levels in the plasma and CSF of AD patients [84–86]. Recenlty, studies have shown that TTR promotes endothelial migration and upregulates angiogenic mediators, including vascular endothelial growth factor, suggesting a pro- angiogenic function under physiological conditions [51]. In contrast, a mutated form of TTR was reported to impair endothelial cell migration, highlighting the importance of structural integrity in TTR-mediated vascular regulation [52]. Interestingly, in pathological contexts such as diabetic retinopathy, TTR appears to exert inhibitory effects on retinal microvascular endothelial cell migration and tube formation, suggesting that its vascular actions may be highly context-dependent [87, 88]. Here, we demonstrate for the first time that TTR is not only expressed in hippocampal capillaries and microglia, but also with a selectively reduced level in both compartments in APP/PS1 mice and AD patients. Remarkably, the decline in capillary TTR levels was also reproduced in a recently reported FMT model, suggesting that the TTR reduction in AD can be recapitulated by changes in gut microbes—alterations frequently reported in aging populations and individuals with neurological disorders [89, 90].

Of note, in this study, the age-dependent decline in TTR expression presented an interesting discrepancy: while we observed reduced TTR in hippocampal capillaries in aged mice, with loss of the dynamic regulation of BBB closure, this pattern was not replicated in human autopsy brain samples. This finding is supported by another study showing variations in TTR levels in CSF across different age groups in healthy controls [85]. Several factors may account for this divergence. First, species differences in lifespan and metabolic rate could accelerate age-related TTR decline in rodents, while humans may maintain compensatory mechanisms that preserve TTR expression in the aging brain. Second, variation in baseline inflammatory states and gut microbiota complexity may shape BBB resilience differently across species. Importantly, this divergence highlights the translational limitation of rodent models while also suggesting that TTR upregulation in humans could represent an adaptive mechanism worth further investigation.

A key finding of this study is that BBB permeability can be bidirectionally regulated by modulating TTR levels, with TTR overexpression not only rescuing LPS-induced hippocampal leakage in young WT mice, but also improving cognitive impairment and reducing neuroinflammation, potentially through alleviating BBB dysfunction in aged WT mice. Unlike humans, aged WT mice rarely develop Aβ plaque deposition. Nevertheless, it is very likely that TTR—including neuron-derived TTR—may mitigate neuroinflammation and cognitive impairment by limiting Aβ accumulation and facilitating its clearance. Further research will focus on the interaction between microglial TTR, neuronal TTR and Aβ aggregation in AD transgenic mice. Notably, TTR is also an amyloid-forming protein which can form fibrils and induce cardiac amyloidosis. The dual role of TTR should be carefully taken into consideration when it is considered as a potential therapeutic target in AD.

## Conclusions

In summary, our study uncovers a previously unrecognized GBA mechanism in which the hippocampal BBB dynamically tightens in response to peripheral inflammation through an age-dependent process driven by TTR. We show that TTR, enriched in hippocampal capillaries and HSM, is indispensable for preserving BBB integrity. Its marked reduction in both AD mice and human patients underscores a pathological loss of this protective mechanism. Together, these findings position TTR as a key molecular link between gut inflammation, BBB regulation, and neurodegeneration—offering a new therapeutic entry point for restoring AD vascular integrity.

## Declarations

### Ethics approval and consent to participate

All human experiments were conducted with approval from the First Affiliated Hospital of Zhejiang University School of Medicine (20240729). The postmortem brain tissues were obtained from the China National Health and Disease Human Brain Tissue Resource Center (Hangzhou, China). All materials were sourced from donors who had provided written informed consent for a brain autopsy and permitted their clinical information for research purposes. All mouse experiments were conducted with approval from the Animal Care and Use Committee of the animal facility at the First Affiliated Hospital of Zhejiang University School of Medicine (20231931).

### Availability of data and materials

The datasets used and/or analysed during the current study are available from the corresponding author on reasonable request. The raw data for scRNA-seq of the DSS model mice hippocampus and the R code necessary to reproduce the analysis and figures in this manuscript have been uploaded to Github at https://github.com/Celbin013/BBB-HSM.

### Funding

National Natural Science Foundation of China under Grant 32530027

Key R&D Program of Zhejiang Province under Grant 2024C03098

Natural Science Foundation of Zhejiang Province under Grant LY24H090006

Innovative Institute of Basic Medical Science of Zhejiang University

### Authors’ contributions

ZX and JYY contributed equally to this work. The order of co–first authors was determined by mutual agreement among the authors.

Conceptualization: ZX, JZ, BX

Investigation: ZX, JYY, JD, YZX, PW, XL, JC, CZ, XDZ

Data analysis: ZX, JYY, YQG Sample collection: ZQF

Visualization: ZX, JYY

Funding acquisition: JZ, BX

Supervision: JZ, BX

Writing – original draft: ZX, JYY

Writing – review & editing: ZX, JYY, BX, JZ

## Acknowledgements

We deeply appreciate the donors for their generous donation of samples. We thank the staff of the Liangzhu Animal Experiment Center at Zhejiang University for their assistance with animal care and the support from the *in vivo* imaging platform. In addition, we thank the staff of the Science Center of the First Affiliated Hospital, Zhejiang University School of Medicine, as well as the imaging platform, for their technical assistance and support.

