## supplementary information for "Transthyretin Contributes to Hippocampal Blood-Brain Barrier Recovery Following Intestinal Inflammation, with Reduced Expression in Alzheimer’s Disease"

**The file includes:**

Figs. S1 to S10

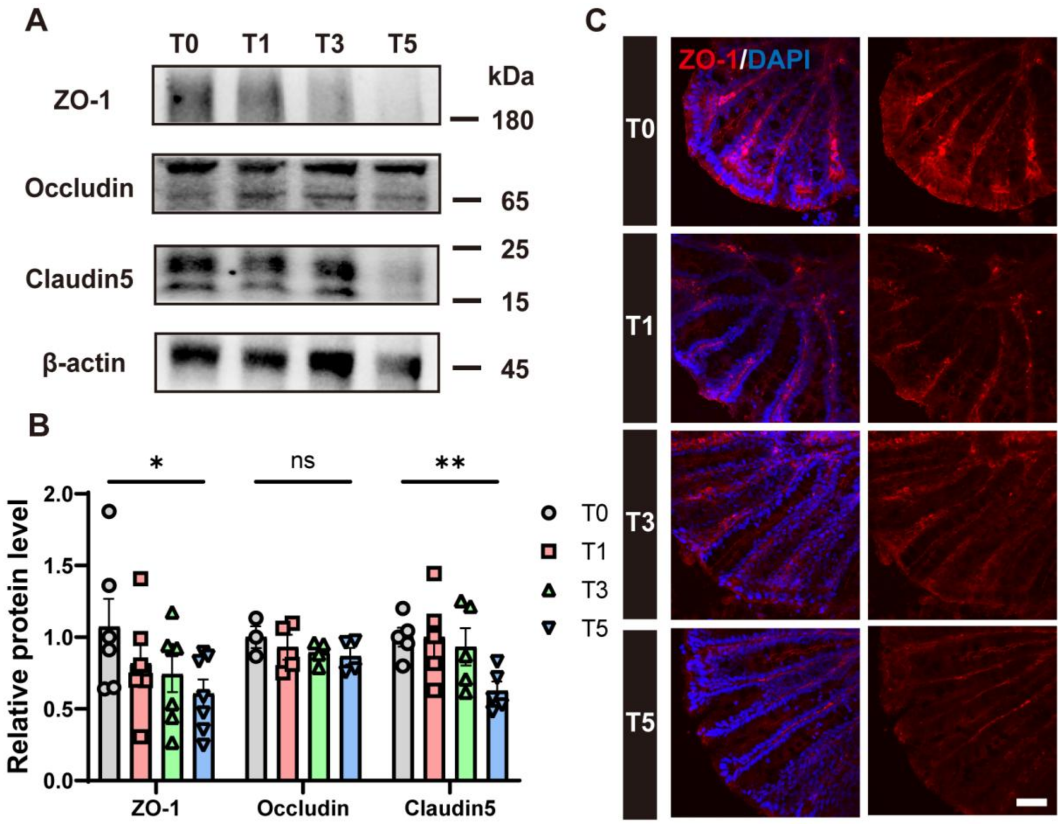


**Supplementary Figure 1 Impaired tight junctions of gut vascular barrier in 2-month-old C57BL/6 mice (2m-WT) during acute colitis. (A-B)** Representative WB images (A) and quantifications (B) of the expression for ZO-1, Occludin and Claudin5 in the colon. For ZO-1, n = 6 mice (T0, T1), n = 7 mice (T3, T5). For Occludin, n = 3 mice (T0), n = 4 mice (T1, T3 and T5). For Claudin5, n = 5 mice for each group. Comparisons between T0 and T5 were performed with unpaired t test. ns, no significance, $*P < 0.05$, $**P < 0.01$. **(C)** Representative z-stack confocal images of ZO-1 (red) in the colon to support the WB quantification. Nuclei were stained with DAPI (blue). Scale bar = 50 μm.

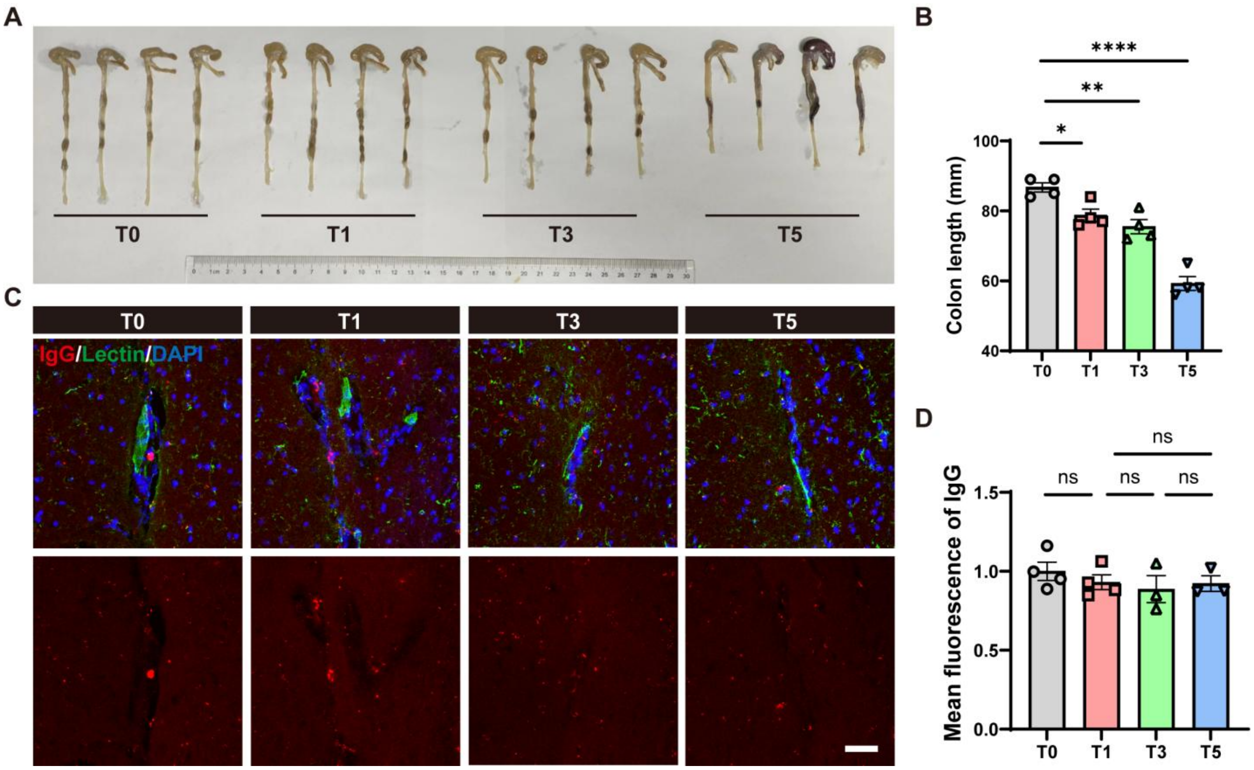


**Supplementary Figure 2 Hippocampal permeability of IgG in 10-month-old C57BL/6 mice (10m-WT) during acute colitis. (A-B)** Image of colon tissues (A) and quantification of the colon length (B). N = 4 mice for each group. **(C-D)** Representative z-stack confocal images (C) and quantification (D) of extravasated plasma IgG (red) around capillaries labeled by Lectin (green) in the Hp. Scale bar = 50 μm. N = 4 mice for T0, T1 and n = 3 mice for T3 and T5. Nuclei were stained with DAPI (blue). Comparisons between groups were performed using one-way ANOVA followed by Tukey's multiple comparisons test. ns, no significance, *$P < 0.05$, **$P < 0.01$, ****$P < 0.0001$.

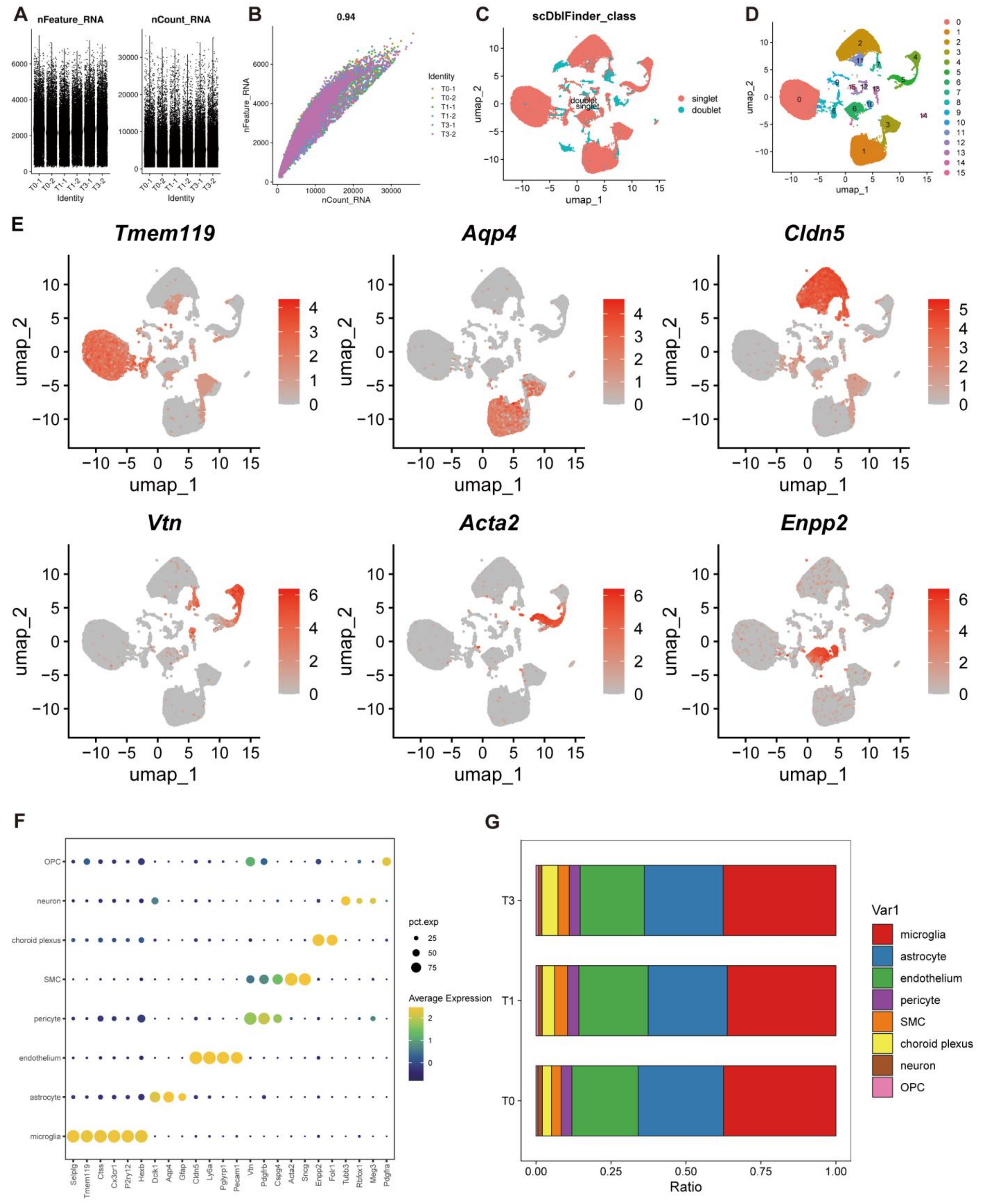


**Supplementary Figure 3 Single-cell RNA sequencing information. (A)** Quantifications of nFeature_RNA and nCount_RNA of each cell in the six samples. **(B)** Linear regression between nFeature_RNA (on the y-axis) and nCount_RNA (on the x-axis) of each cell in the six samples.

R = 0.94. **(C)** UMAP plot showing doublet removed by scDblFinder. **(D)** UMAP plot of cells in the Hp. Different time points are labeled in different colors, and no obvious distribution differences are observed among the three groups. Sample size: T0-1, 11,335 cells; T0-2, 11,344 cells; T1-1, 11,478 cells; T1-2, 11,724 cells; T3-1, 11,026 cells; T3-2, 10,515 cells. Each sample consists of the bilateral hippocampi pooled from three mice. **(E)** UMAP plots showing expression of representative markers used to annotate microglia (*Tmem119*), astrocyte (*Aqp4*), endothelium (*Cldn5*), pericyte (*Vtn*), SMC (*Acta2*) and choroid plexus (*Enpp2*). **(F)** Dotplot showing representative markers used to annotate each cellular population. Dot size represented the percentage expression (pct.exp) of the gene in each cell type. Dot color represented average expression value of each gene in each cell type. **(G)** Cell ratio of microglia, astrocyte, endothelium, pericyte, SMC, choroid plexus, neuron and OPC at T0, T1 and T3.

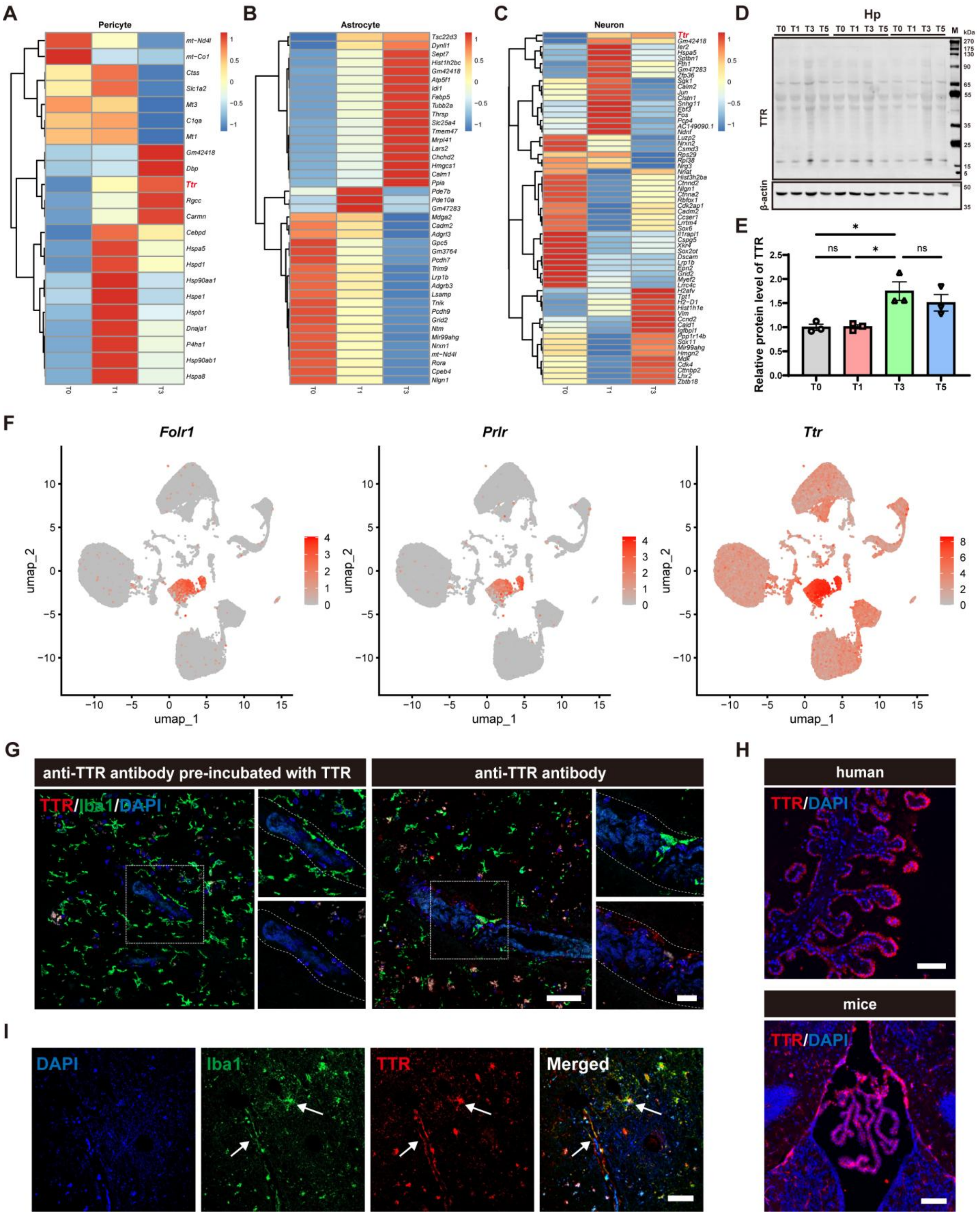


**Supplementary Figure 4 Supplementary DEGs heatmaps related to Fig. 2 and TTR expression validation. (A-C)** Heatmap showing the DEGs between T0, T1 and T3 in pericyte (A), astrocyte (B) and neuron (C). **(D-E)** Representative WB images (D) and quantification (E)

of TTR expression level in the Hp of 2m-WT. N = 3 mice for each group. Comparisons between groups were performed using one-way ANOVA followed by Tukey's multiple comparisons test. ns, no significance, *$P < 0.05$. **(F)** UMAP plots showing expression of *Folr1*, *Prlr* and *Ttr* in all cell types. **(G)** Representative z-stack confocal images of TTR (red) and microglia labeled by Iba1 (green) around human hippocampal capillaries. Anti-TTR antibody pre-incubated with TTR protein was used to validate the specificity of the antibody. Nuclei were stained with DAPI (blue). Scale bar = 100 µm (left) and 20 µm (right). **(H)** Representative IF images showing strong signals of TTR in the choroid plexus of both human and mouse brain tissues. Scale bar = 100 µm. **(I)** Representative 2D confocal images showing co-localization of TTR (red) and microglia labeled by Iba1 (green) in human post-mortem tissue. Scale bar = 50 µm.

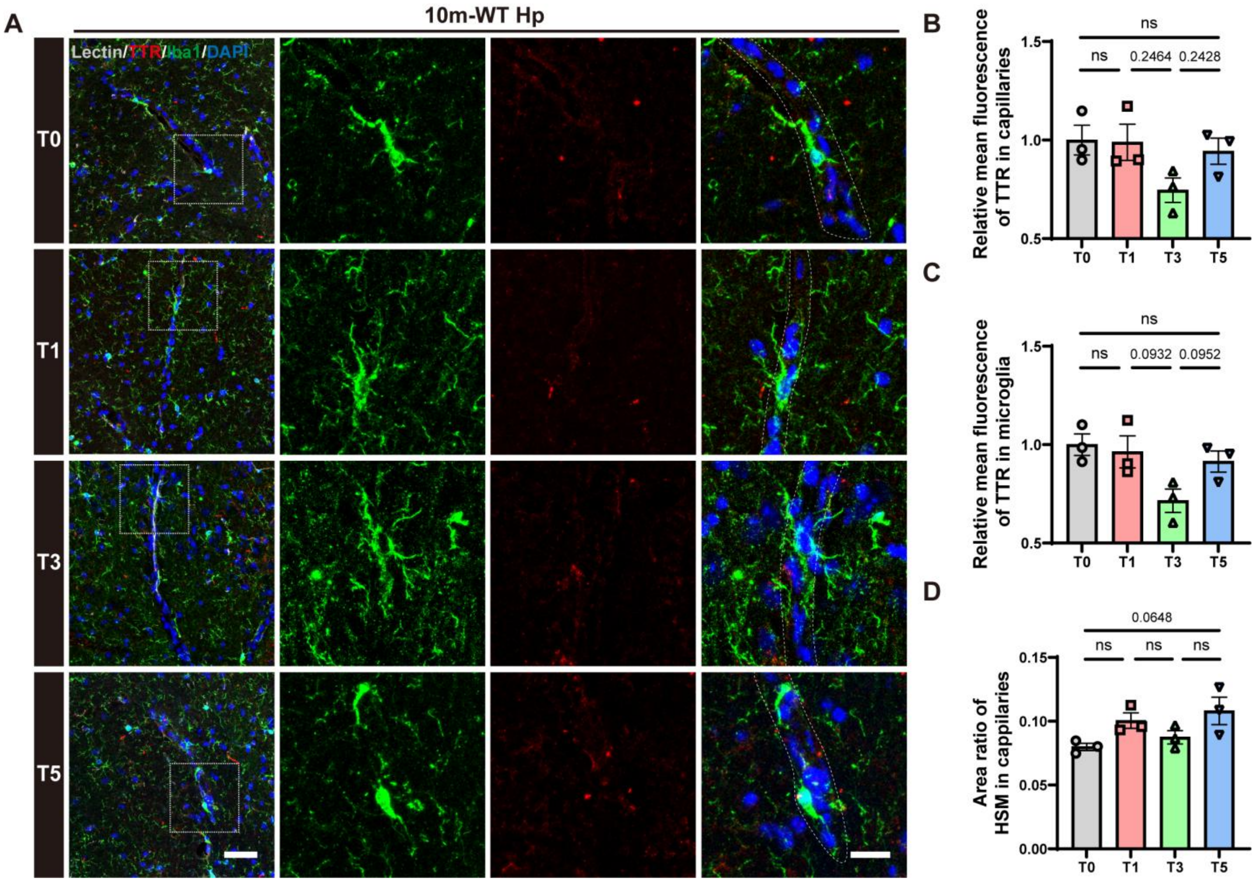


**Supplementary Figure 5 Hippocampal staining of capillaries, Iba1 and TTR in 10m-WT during acute colitis.** **(A)** Representative confocal images of TTR (red) and microglia labeled by Iba1 (green) around capillaries labeled by Lectin (grey). The merged images in the fourth column display signals in the absence of Lectin. Scale bar = 50 μm (left) and 20 μm (right). **(B)** Quantification of TTR level in the hippocampal capillaries. **(C)** Quantification of TTR level in HSM. **(D)** Quantification of area ratio of HSM in hippocampal capillaries. Nuclei were stained with DAPI (blue). N = 3 mice for each group. Comparisons between groups were performed using one-way ANOVA followed by Tukey's multiple comparisons test. ns, no significance.

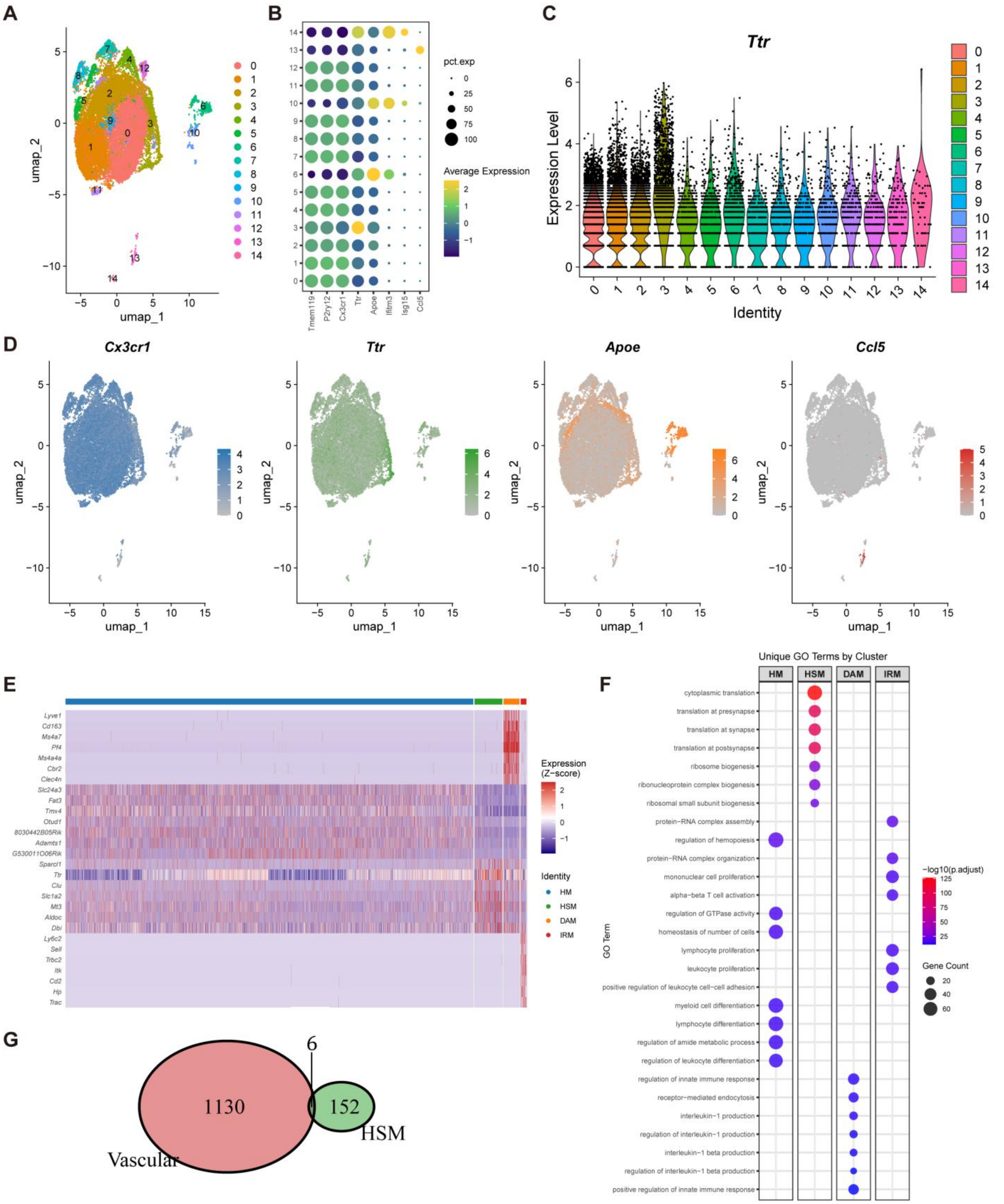


**Supplementary Figure 6 Microglial subtypes information related to Fig. 2. (A)** UMAP plots showing 16 microglia subclusters. **(B)** Dotplot showing representative markers used to annotate microglial subtypes. Dot size represented the percentage expression (pct.exp) of the gene. Dot color represented the average expression value of each gene. **(C)** Vlnplot showing *Ttr* expression in all microglia subclusters. **(D)** UMAP plots showing expression of representative markers used to annotate homeostatic microglia (HM, *Cx3xr1*), high-sensitive microglia (HSM, *Ttr*), disease-associated microglia (DAM, *Apoe*) and interferon-responsive microglia (IRM, *Ccl5*). **(E)** Heatmap showing the top 7 DEGs among the four microglial subtypes. **(F)** Dotplot showing top 7 unique gene ontology (GO) terms in the four microglial subtypes. Dot size represented the gene count. Dot color represented the significance of pathway enrichment (-log10(p.adjust)). **(G)** Venn plot showing overlap of vascular gene sets and differentially expressed genes (DEGs) in HSM.

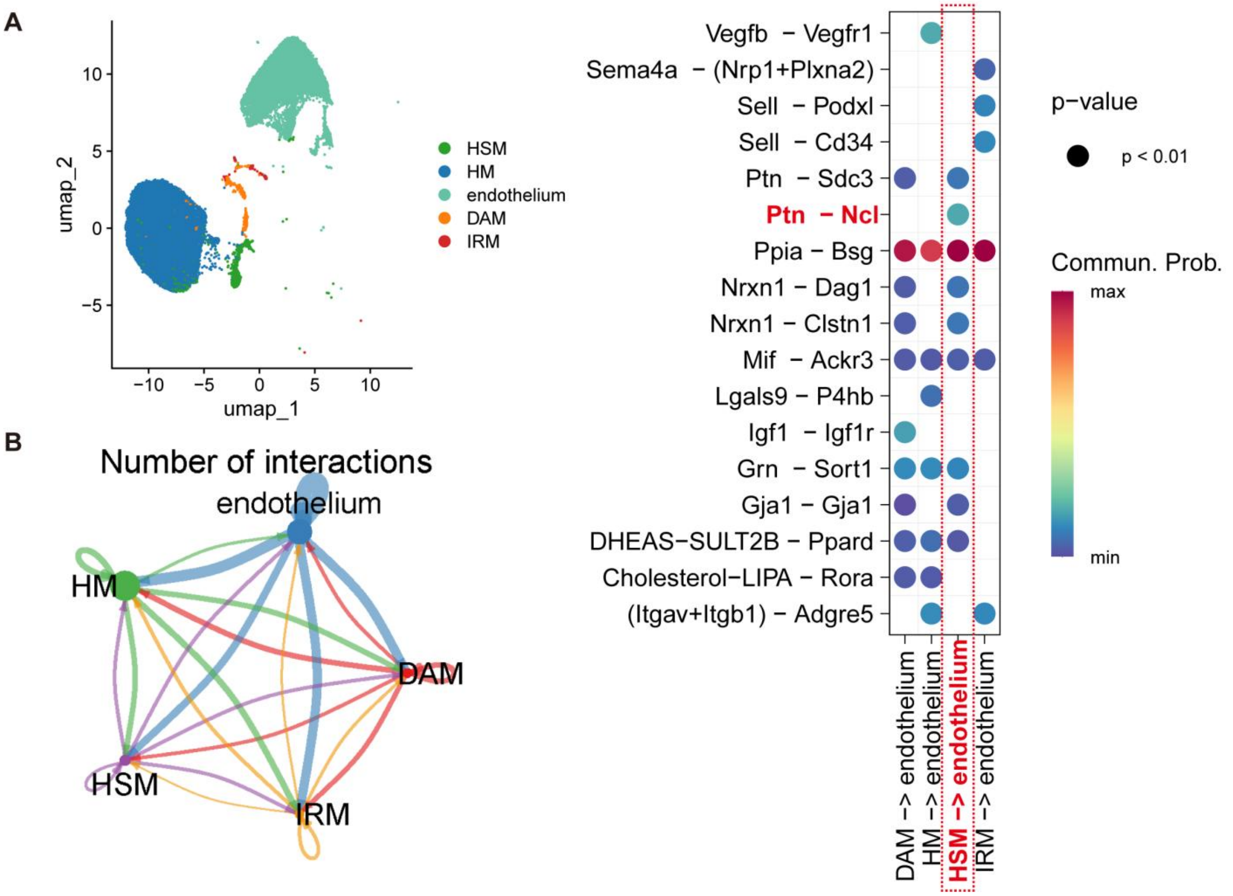


**Supplementary Figure 7 Interactions network between microglial subtypes and endothelium. (A)** UMAP plot showing the distribution of HM, DAM, IRM, HSM and endothelium. **(B)** Circle plots illustrating the interaction network among HM, DAM, IRM, HSM and endothelium. The width of the lines connecting these cell types represents the number of interactions. **(C)** Bubble plot illustrating gene interactions among HM, DAM, IRM, HSM and endothelium. The x axe represents transitions between different cell types and the y axe represents different pairs of interacting genes. The size of the bubbles denotes the statistical significance of the interactions and the color of the bubbles corresponds to the strength of the interactions.

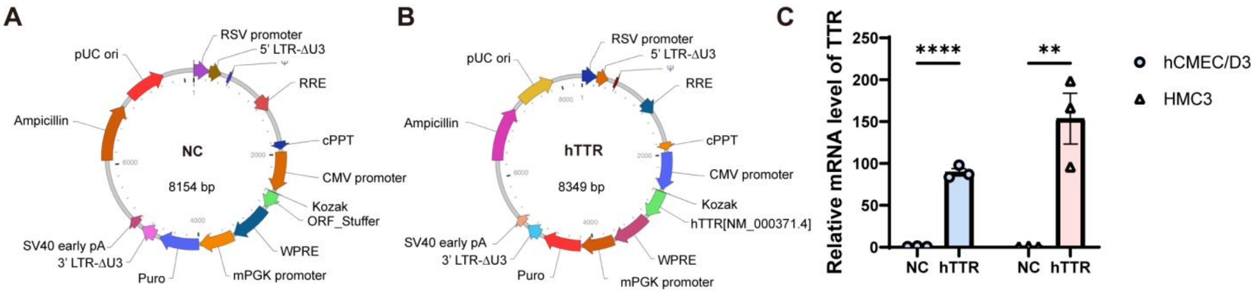


**Supplementary Figure 8 Overexpression of TTR in hCMEC/D3 and HMC3. (A-B)** Atlas of hTTR expressing lentivirus (B) and negative lentivirus vectors (A). **(C)** Quantifications of *TTR* mRNA levels in hCMEC/D3 and HMC3 infected with hTTR-expressing lentivirus (hTTR) and negative lentivirus vectors (NC). N = 3 wells. Comparisons between groups were performed with unpaired t test. $^{**}P < 0.01$, $^{****}P < 0.0001$.

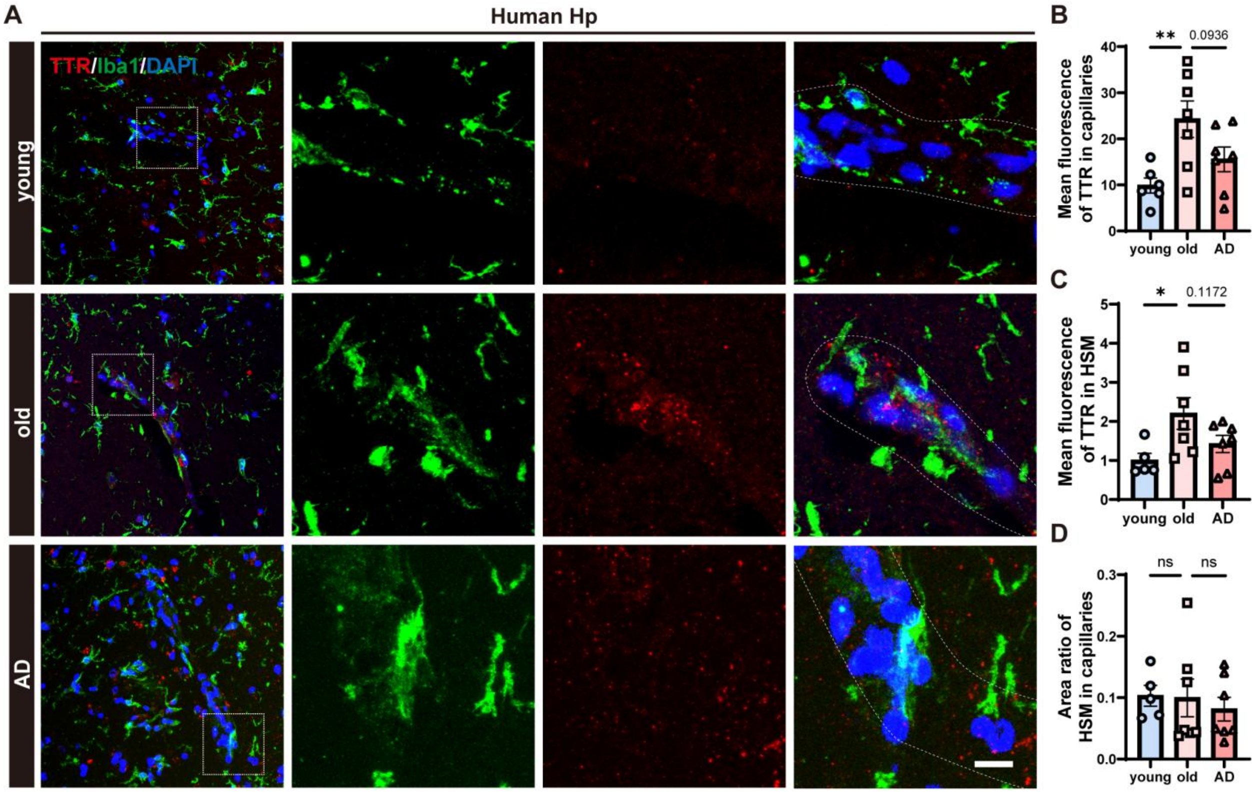


**Supplementary Figure 9 TTR expression in hippocampal capillaries decreases in AD patients. (A)** Representative z-stack confocal images of TTR (red) and microglia labeled by Iba1 (green) around human hippocampal capillaries. Scale bar = 50 μm. **(B-C)** Quantification of TTR in hippocampal capillaries (B) and HSM (C) of young healthy control (young), old healthy control (old) and AD patients (AD). **(D)** Quantification of the area ratio of HSM in hippocampal capillaries of the three groups. N = 6 donors for young and n = 7 donors for old and AD. Nuclei were stained with DAPI (blue). Comparisons between groups were performed with unpaired t test. ns, no significance, *$P < 0.05$, **$P < 0.01$.

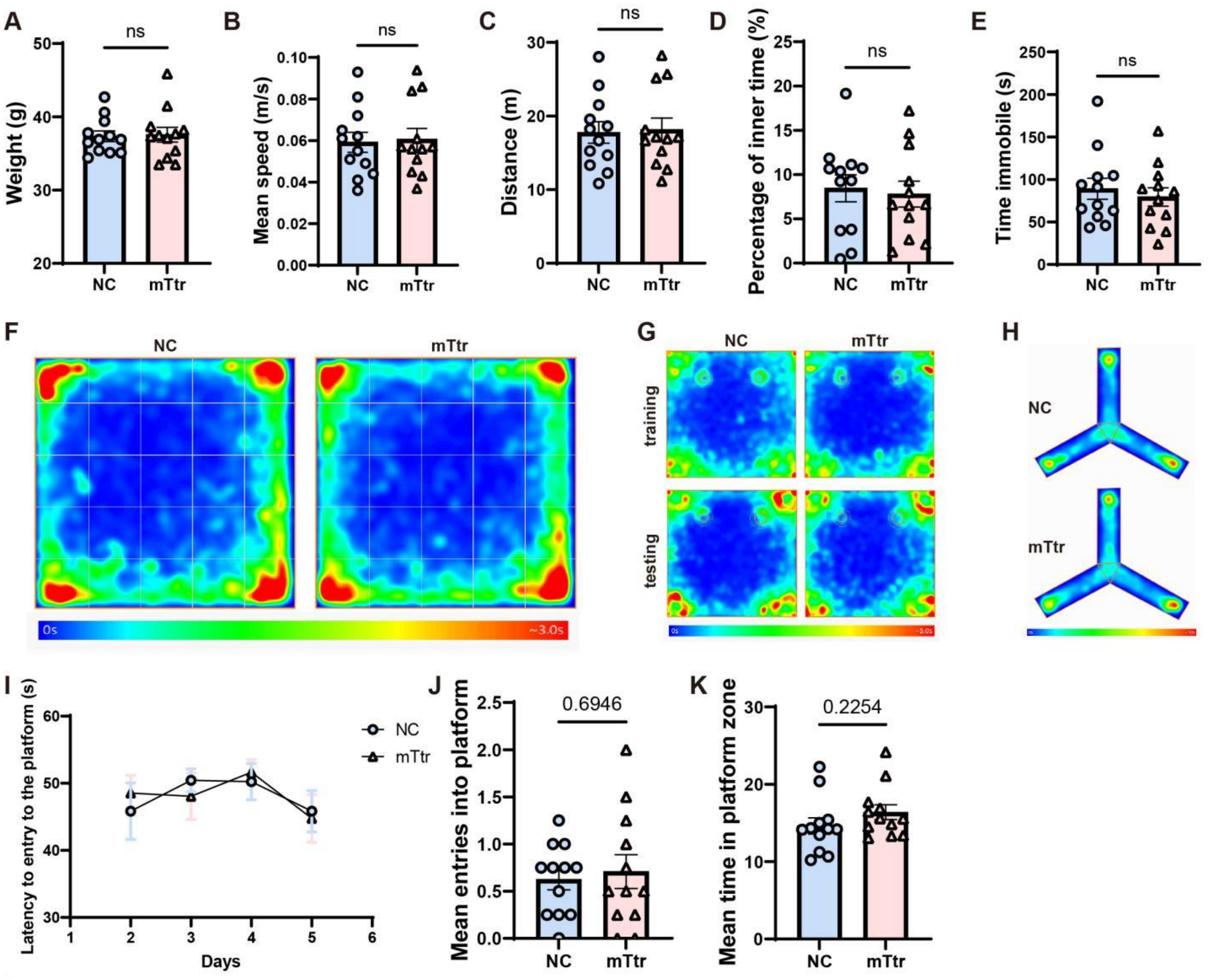


**Supplementary Figure 10 Open filed test and Morris water maze in NC and mTtr. (A-E)** Quantifications of weight (A), mean speed (B), distance (C), percentage of inner time (D) and time immobile (E) of 10-month-old NC and mTtr. **(F-H)** Heatmap showing routes of all mice in the same group in Open field test (F), Novel object recognition (G) and Y-maze (H). **(I)** Quantification of latency to entry to the platform on Days 2-5. **(J-K)** Quantification of mean entries into platform (J) and mean time in platform zone (K) on Day 6. N = 12 mice for each group. Comparisons between groups were performed with unpaired t test. ns, no significance.
